# A Continuum of Atrial Peristalsis Initiates the Bicuspid to Quadricuspid Valve Transition

**DOI:** 10.64898/2026.05.04.722768

**Authors:** Jing Wang, Aaron L. Brown, Peng Zhao, Charlie Z. Zheng, Jau-Nian Chen, Bin Zhou, Jiandong Liu, Tomohiro Yokota, Alexander D. Kaiser, Seul-Ki Park, Alison L. Marsden, Tzung K. Hsiai

**Affiliations:** Department of Bioengineering, UCLA, Los Angeles, CA, USA; Department of Mechanical Engineering, Stanford University, Stanford, CA, USA; Division of Cardiology, Department of Medicine, School of Medicine, UCLA, Los Angeles, CA, USA; Department of Medicine, Greater Los Angeles VA Healthcare System, Los Angeles, CA, USA; Department of Molecular, Cell, and Developmental Biology, UCLA, Los Angeles, CA, USA; School of Medicine, Stanford University, Stanford, CA, USA; Department of Pediatrics, University of Chicago, IL, USA; Department of Pathology and Laboratory Medicine, School of Medicine, University of North Carolina at Chapel Hill, NC, USA; Department of Cardiothoracic Surgery, Stanford University, Stanford, CA, USA; Departments of Pediatrics and Bioengineering, Stanford University, Stanford, CA, USA

## Abstract

How chamber-specific hemodynamic shear forces coordinate the spatial and temporal patterning of cardiac valves remains poorly understood. The zebrafish atrioventricular (AV) valve transitions from 2-cusp to 4-cusp at 14 days post-fertilization (dpf) through commissural remodeling, differential cell proliferation, and chondrogenic matrix secretion. Orthogonal valvular axes are present at 25 hpf (hours), preceding endocardial cushion formation. AV canal remodeling aligns the shear stress with leaflet formation at 48 hpf and 14 dpf. At 28 hpf, the ventricle develops functional syncytium, while the atrial peristalsis remains and is necessary for the Klf2a-Snai1b signaling in Notch-negative and *ccm1*-positive endocardial cells. Without atrial contraction, the 2-to 4-cusp transition (86.7%) fails to occur. In the absence of ventricular contraction and relaxation, the formation of the first pair of leaflets (100%) does not initiate. As a corollary, we demonstrate that KLF4/SNAI1 expression at the commissure is conserved in the mammalian AV 2- to 4-cushion transition.

## Introduction

Congenital bicuspid valvular heart disease poses a considerable healthcare burden on the aging societies worldwide ^1^. The mammalian heart is equipped with four sets of valves: aortic, pulmonary, mitral, and tricuspid. Each has a unique cellular and extracellular matrix (ECM) architecture, encompassing a myriad of molecular pathways that may contribute to stenosis or regurgitation in a diseased valve ^2^. Efforts to identify therapeutic targets are complicated by an incomplete understanding of how congenital valvular structures originate and interact with their biomechanical environments ^3^. For example, patients with a bicuspid aortic valve (BAV) are prone to accelerated progression of calcification, aortic stenosis, and aortic aneurysm ^4^. It is postulated that BAV leads to locally disturbed hemodynamics, which prime osteogenic pathways in valvular interstitial cells (VICs) via mechanical transducers (e.g., Piezo1, Notch) and trigger calcium deposition ^5–8^ Nevertheless, it remains unclear how mechanical forces may have contributed to the abnormal number of leaflet during valvular formation.

To address this knowledge gap, substantial research has focused on the mechanobiology of valvular formation across vertebrate models ^9–15^. Studies in murine and avian models have elucidated the fundamental processes of valvular development. The aortic/pulmonary and mitral/tricuspid valves are derived from the outflow tract (OFT) and atrioventricular (AV) cushions, respectively. The OFT and atrioventricular canal (AVC) initially consist of one pair of “major” cushions, which later develop into another pair of “lateral (or intercalated)” cushions. The major cushions will fuse as the embryonic heart undergoes septation, which divides the left and right atria/ventricles, and the aortic and pulmonary outflow tracts.

Meanwhile, the four cushions reorganize into the primordial leaflets. During initiation, the cushions are composed of acellular ECM (cardiac jelly) secreted by the myocardium. Hemodynamic shear stress-responsive signaling, including KLF2/4, WNT, NOTCH1, and NFATc1, coordinates the epithelial-mesenchymal transition (EMT) in endocardial cells (also known as EndoMT) ^9,17,18^. Activated endocardial cells express mesenchymal transcription factors, such as *Snai1*, *Snai2 (Slug)*, and *Twist1/2*, as they invade the underlying cardiac jelly and differentiate into valve interstitial cells (VICs) ^19–22^. Other VIC lineages also contribute to the population, depending on the particular cushion, including neural crest, epicardium, and the second heart field (SHF). VICs remodel and stratify the ECM, a process that continues postnatally until adulthood ^23,24^.

Zebrafish has provided critical insights into the mechanobiology of cardiac development due to its accessibility for imaging and genetic modifications ^12,25^. The embryo is transparent and readily supports fluorescent reporters, enabling longitudinal tracking of cardiac morphogenesis at single-cell resolution. The zebrafish AV valve shares a flow-mediated initiation with the mammalian EndoMT at 50 hours post-fertilization (hpf), in which Notch-positive (^+^) endocardial cells at the AV cushion secrete Wnt9a via *Klf2a* activation, triggering the ingression of adjacent Notch-negative/Klf2a-low endocardial cells ^26^. These cells proliferate to form a pocket within the cushion, which unfolds at 65-80 hpf to give rise to primordial leaflets. Nfatc1 modulates this step by regulating *twist1b* expression, and loss of Nfatc1 impairs VIC recruitment, leading to hypocellular, insufficient valves.

Computational fluid dynamics (CFD) simulations have revealed that high oscillatory shear activates the convergent movement of AVC endothelium ^28^. Klf2a activates the inhibitor gene, *ccm1* (*krit1*), which dampens Klf2a’s effect and restricts it to regions of high shear ^29^. In *ccm1* mutants, both *klf2a* and *notch1b* lose their AVC localization, and endocardial cells fail to ingress. In OFT valves, increased hemodynamic shear stress from isoproterenol treatment (contractility) or EPO mRNA injection (viscosity) augments the leaflet volume and the number of VICs ^30^. Despite the identification of additional mechanosensitive pathways ^31–33^, most research focuses on the embryonic phase, when the AV valve is bicuspid. A recent study uncovers the quadricuspid AV valve in adult zebrafish, and the number of cusps (leaflets) is influenced by intracardiac flow ^34^. This suggests that the zebrafish AV valve undergoes a 2- to 4-cusp transition similar to that of mammalian AV and OFT cushions. Nevertheless, it remains unclear how hemodynamic shear stress coordinates the location and timing of leaflet initiation.

In this study, we reveal the patterning of leaflets via chamber-specific shear stress in the zebrafish AV valves, which undergo the 2- to 4-cusp transition at 14 days post-fertilization (dpf) through commissural remodeling and flow-mediated endocardial differentiation. From 15 to 28 dpf, the lateral cushions are remodeled into semilunar cusps and establish themselves in the AVC via differential VIC proliferation relative to the pre-existing cusps. From 28 to 45 dpf, laser microdissection-assisted RNA sequencing and spatial RNA sequencing suggest that first the pre-existing cusps, then the lateral cusps, mature by upregulating valve-specific genes related to chondrogenesis, retinoic acid signaling, and glycogen metabolism. More importantly, we demonstrate that the orthogonal valvular axes for the first pair of leaflets (superior and inferior cusps) are established as early as 25 hpf, long before endocardial cushions are present. Prior to the initiation of each cusp pair (at 48 hpf and 14 dpf), the AVC undergoes remodeling that redistributes the shear stress along the left and right lateral valvular axes.

Furthermore, we characterize the distinct mode of contraction between the ventricle (synchronous/syncytial) and the atrium (peristaltic) from 28 to 33 hpf. Mutants of chamber-specific myosin heavy chains (*haf* and *wea*) reveal that loss of atrial peristalsis disrupts the shear stress along the commissure of 2-cusp AV valves, impairing Klf2a-Snai1b signaling in the Notch-negative/*ccm1*-positive endocardial cells. In 86.7% of *wea* mutants (absence of atrial contraction), the second pair of cusps fails to develop, while in 100% of the *haf* mutants, the first pair of cusps fails to develop. As a corollary, we demonstrate the conserved expression of KLF4 and SNAI1 in murine OFT and AVC during the 2-cushion to 4-cushion transition. Thus, the continuum of atrial peristalsis generates a shear stress gradient to coordinate the formation of the first pair of leaflets and initiation of the second pair during AV valvulogenesis via Klf2a-Snai1b signaling in the remodeled commissure, where the shear-responsive Notch-negative and *ccm1*-positive endocardial cells are implicated in initiating the 2-cusp to 4-cusp AV transition.

## Results

### Temporal dynamics of atrioventricular valve undergoing 2- to 4-cusp transition

From 14 to 45 dpf, the Notch signaling reporter line *Tg(TP1:EGFP)* ^35^ labels the free edges of the zebrafish atrioventricular (AV) valve, which transitions from 2-cusp to 4-cusp: superior (SUP), inferior (INF), left lateral (LL), and right lateral (RL) (**Figure 1a-b**). EdU (5-ethynyl-2′-deoxyuridine) labeling from 14 to 15 dpf reveals that the valvular commissures remodel via endocardial proliferation, separating the SUP/INF leaflets before new valvular interstitial cells (VICs) appear (**Figure 1c**). Proliferating VICs form the initial endocardial cushion that protrudes into the AV canal (AVC). The cushion then delaminates from the AVC wall to form the early LL/RL cusps with a luminal space (**Supplementary Figure 1a**), as described in the formation of embryonic SUP/INF cusps . Finally, the lateral cusps develop from monolayer to multilayer VICs during maturation.

**Figure 1.**
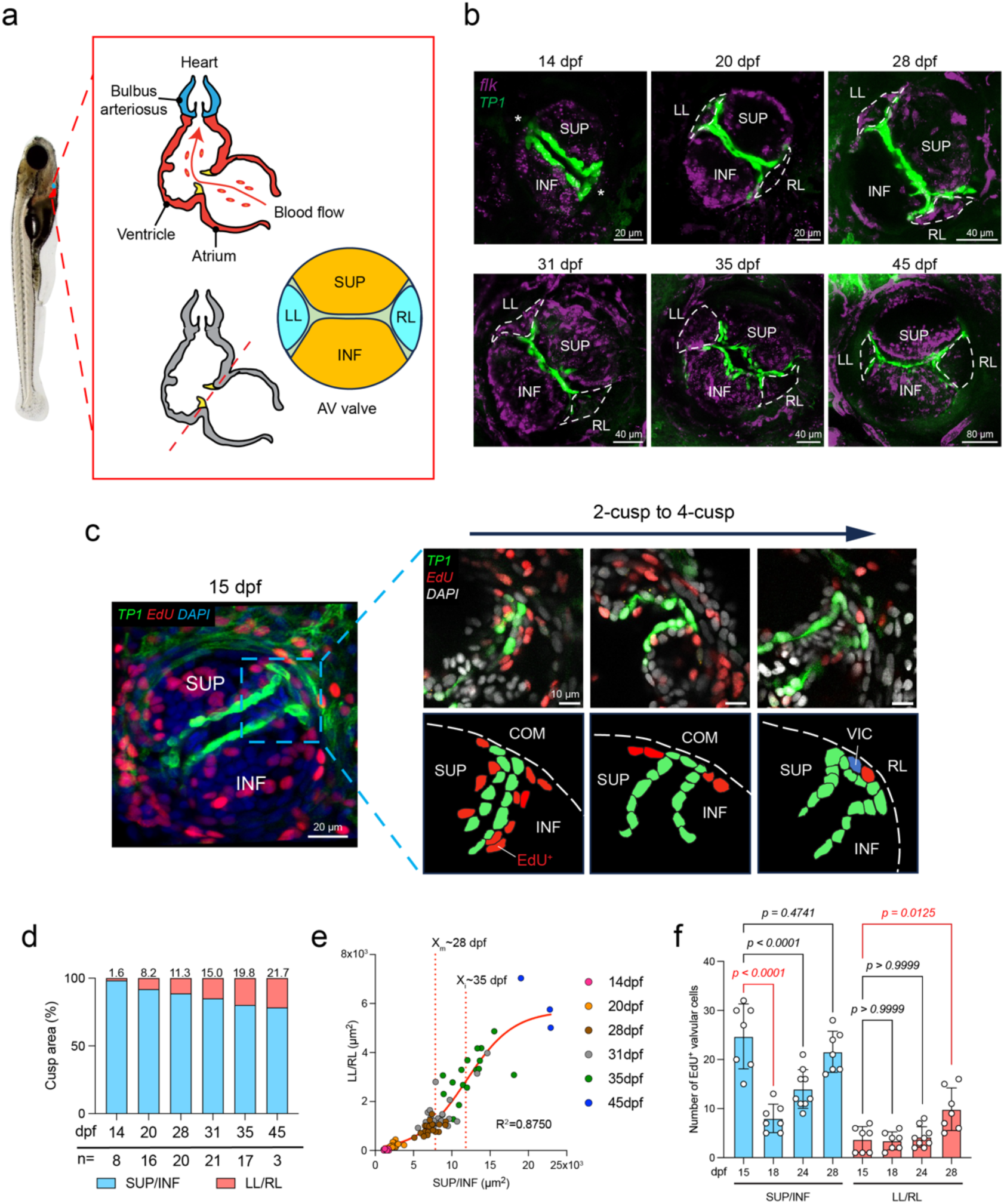
A 2- to 4-cusp AV valve transition develops between 14 and 45 days post-fertilization (dpf). **(a)** A zebrafish larva (left panel) at 12 dpf is shown, with its heart highlighted. The right panel illustrates the two-chambered cardiac anatomy and the quadricuspid atrioventricular (AV) valve. **(b)** From 14 to 45 dpf, confocal imaging of hearts in *Tg(TP1:EGFP; flk:mCherry)* larvae during the transition from 2-cusp to 4-cusp. Dashed lines indicate the developing lateral cusps extending from the commissures (asterisks). **(c)** At 15 dpf, confocal imaging of EdU-labeled hearts (24-hour incubation) from *Tg(TP1:EGFP)* larvae (n = 7) shows that the commissure remodels through endothelial proliferation (highlighted in the blue box) before VIC differentiation, after which VICs proliferate to initiate lateral cusp formation. **(d)** From 14 to 45 dpf, the average area of the left/right lateral cusps (LL/RL) increases more rapidly than that of the pre-existing superior/inferior cusps (SUP/INF), as indicated by the increasing percentage over time. The number (n) of hearts analyzed at each time point is displayed. Source data are available in a Source Data file. **(e)** From 14 to 45 days post-fertilization (dpf), the change in the average area of left/right lateral (LL/RL) versus pre-existing superior/inferior (SUP/INF) cusps follows a logistic growth curve (refer to the Method section for the equation). The relative growth rate of lateral cusps peaked at approximately **35 dpf** (X_i_ = 11.9x10^3^ µm^2^), while the period of maximal growth acceleration occurred earlier at **∼28 dpf** (X_m_ = 7.84x10^3^ µm^2^), identifying the inflection point of greatest augmentation in the second derivative. **(f)** The total number of EdU^+^ valvular cells in the pre-existing superior/inferior cusps (SUP/INF) declines significantly from 15 days post-fertilization (dpf) to 18 dpf, and this number does not recover until 28 dpf. The cell count in the left/right lateral cusps (LL/RL) remains relatively constant from 15 dpf to 24 dpf but increases significantly at 28 dpf. All values are presented as mean ± standard deviation (SD). The p-value is indicated for each comparison. The number of hearts analyzed was n = 7 for 15, 18, and 28 dpf, and n = 9 for 24 dpf. An ordinary two-way ANOVA followed by Sidak’s multiple-comparisons test was used to assess statistical significance. Source data are provided in a Source Data file. Anatomic labels: SUP, superior cusp; INF, inferior cusp; LL, left lateral cusp; RL, right lateral cusp; COM, commissure; VIC, valve interstitial cell.

The average area of the LL/RL cusps increases faster than that of the SUP/INF cusps, rising from 1.6% of total cusp area at 14 dpf to 21.7% at 45 dpf (**Figure 1d**). The change in lateral vs. pre-existing cusp area over time follows a logistic growth curve (R^2^=0.8750; equation in Method), indicating that the growth rate of lateral cusps (relative to the pre-existing cusps) also increases over time, reaching a maximum at the “inflection point” around 35 dpf (**Figure 1e**). The most significant increase in relative growth rate (i.e., the maximum of the second derivative) occurs around 28 dpf.

In SUP/INF cusps, EdU assays show a significant decrease in the number of proliferating cells from 15 dpf (24.7 cells/heart) to 18 dpf (8 cells/heart, *p* < 0.0001) and 24 dpf (14 cells/heart, *p* < 0.0001), with no recovery until 28 dpf (21.6 cells/heart, *p* = 0.4741, n = 7 for 15/18/28 dpf, n = 9 for 24 dpf) (**Figure 1f**). In contrast, the number of EdU^+^ cells in LL/RL cusps remained similar until a significant increase at 28 dpf (9.9 cells/heart, *p* = 0.0125 vs. 3.7 cells/heart at 15 dpf). The dynamics of cell proliferation align with those of the cusp area.

To elucidate the underlying transcriptomic landscape, we perform laser microdissection-assisted RNA sequencing of AV valve tissue from 28, 35, and 45 dpf (**Figure 2a**, **Supplementary Figure 1b;** n = 3 per time point). Before 28 dpf, the AV valves were too small for precise dissection. From 28 dpf to 45 dpf, the sequencing demonstrates significantly reduced expression of cell proliferation markers *pcna* (*p-adjust* = 0.00589) and *mki67* (*p-adjust* = 0.00113) (**Supplementary Figure 1c-e**). However, from 28 dpf to 35 and/or 45 dpf, multiple chondrogenic genes are up-regulated, including a recently discovered cartilage-specific proteoglycan gene, *snorc* ^37^ (**Supplementary Figure 2b,** *p-adjust* = 0.01800 vs. 35 dpf, *p-adjust* = 0.00044 vs. 45 dpf). The sequencing results are corroborated by whole-mount *in situ* hybridization of *snorc* mRNA in the *Tg(TP1:EGFP)* hearts (20/28/35 dpf), in which the increase in *snorc* is more prominent in SUP/INF cusps than in LL/RL cusps (**Supplementary Figure 2a,c**). In the lateral cusps, the expression of *snorc* does not become statistically different until 35 dpf (*p* = 0.0395 vs. 20 dpf, n = 8 for 20/28 dpf, n = 6 for 35 dpf), suggesting slower ECM maturation. The remaining chondrogenic genes encode matrix metalloproteinase (*mmp2*), proteoglycan (*prg4*), glycoprotein (*pcolce2a*), collagen (*col2a1b*), and cytokeratin (*krt4*, *krt91*) (**Supplementary Figure 2d**). In parallel, the sequencing analysis shows upregulation of genes involved in ciliation (*ttc9c*, *tbc1d31*), protein synthesis (*eef1a1a*), retinoic acid signaling (*rbp4*, *abca3b*, *greb1l*), and glycogen metabolism (*gyg1b*, *pygl*, *ppp1r3aa*) (**Supplementary Figure 3**). A recently published Stereo-seq dataset of adult zebrafish hearts ^38^ confirms that 11 of the 16 above-described genes are clearly enriched in the AV valve (**Figure 2b-e**).

**Figure 2.**
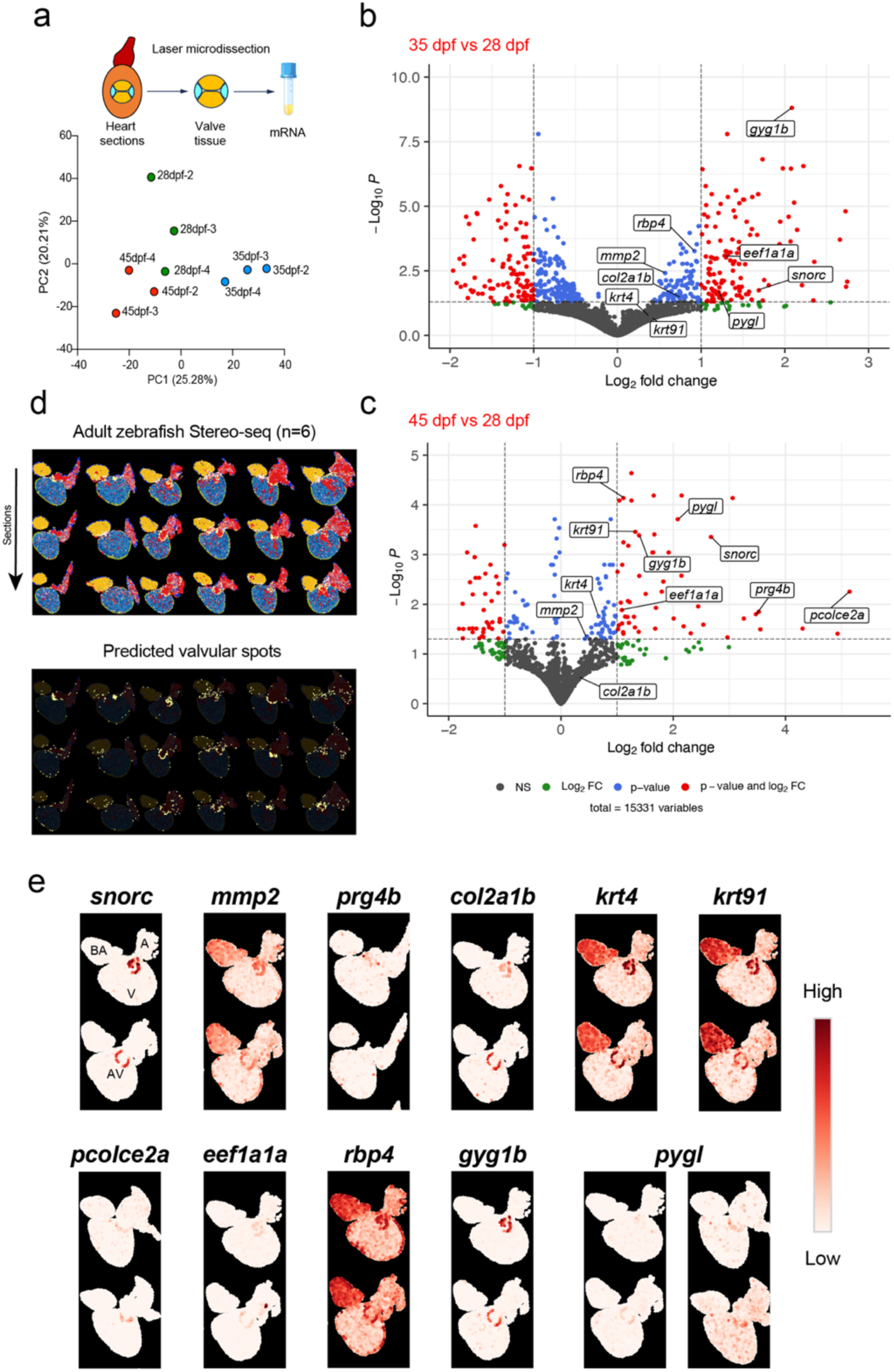
Laser microdissection-assisted RNA sequencing reveals valvular maturation programs following the 2- to 4-cusp transition. **(a)** Laser microdissection-assisted RNA sequencing is performed on AV valve tissue from 28-, 35-, and 45-dpf WT zebrafish. Four samples from each time point are collected for sequencing, each containing 30∼60 tissue pieces (depending on age) cut from ∼10 hearts. One outlier from each time point is discarded based on the principal component (PC) analysis plot of normalized gene counts. Percentages of variance explained by the first two PCs are indicated on each axis. Source data are provided as a Source Data file. **(b - c)** Volcano plots show the log-fold changes and Benjamini & Hochberg (1995) adjusted *p*-values for upregulated genes (labeled with lines) at 35 or 45 dpf versus 28 dpf. The genes shown here are validated in the spatial transcriptomic atlas, involved in chondrogenesis (snorc, *mmp2, prg4b, col2a1b, krt4, krt91, pcolce2a*), protein synthesis (*eef1a1a*), retinoic acid signaling (*rbp4*), and glycogen metabolism (*gyg1b*, *pygl*). Original *p*-values are computed using the Wald test in DESeq2. Dashed lines mark the selected thresholds for log-fold changes (-1/1) and adjusted *p*-values (0.05). Colors indicate genes that pass the thresholds for log-fold changes (green), adjusted *p*-values (blue), or both (red). Normalized gene counts and adjusted *p*-values are shown in Supplementary Figures 2 & 3. **(d)** Spatial clustering of major cell-type domains (colored spots) in adult WT zebrafish heart sections from a published Stereo-seq dataset. The right panel shows the authors’ predicted valvular domain (yellow spots, marker genes: *angptl7* and *abi3bpb*). The dataset includes six hearts (3 sections each). **(e)** Differentially expressed genes identified by laser microdissection-assisted RNA sequencing show spatial enrichment within valvular domains. The color scale indicates expression levels. Anatomic labels: V, ventricle; A, atrium; AV, atrioventricular valve; BA, bulbus arteriosus.

Overall, our data demonstrates that the lateral cusps of the zebrafish AV valve arise from remodeled commissures at 14 dpf and undergo proliferation and delamination similar to the pre-existing cusps. The LL/RL cusps exhibit faster growth in cusp area until 35 dpf, driven by reduced proliferation in the SUP/INF cusps between 15 dpf and 18 dpf. Beyond 35 dpf, the AV valve initiates its maturation program in the SUP/INF cusps before the LL/RL cusps.

### Valvular axes during ventricular peristaltic-to-synchronous contraction

To clarify how hemodynamic shear stress may pattern the two pairs of leaflets within the AVC, we first examined the establishment of valvular axes in *Tg(TP1:EGFP)* embryos throughout cardiac development. At 25 hpf, the linear heart tube exhibits two orthogonal axes within its elliptical cross-section: the superior-inferior (S-I) long axis, along which the SUP/INF cusps will develop, and the left-right (L-R) short axis for the LL/RL cusps (**Figure 3a, Supplementary Video 1**). From 25 hpf to 72 hpf, the spatial orientation of these axes is conserved, although at 48 hpf the heart undergoes looping and ballooning (**Supplementary Figure 4a, Supplementary Video 3**). From 28 hpf to 33 hpf, peristaltic ventricular contraction becomes synchronous (i.e., a functional syncytium), as indicated by the slowing and eventual cessation of the peristaltic wave at the AVC (**Supplementary Figure 4b**). At 54 hpf, atrial contraction remains peristaltic during SUP/INF cusp initiation, as shown by the *Tg(fli1a:LifeAct-GFP)* endocardial reporter (**Supplementary Figure 4c, Supplementary Video 4**).

**Figure 3.**
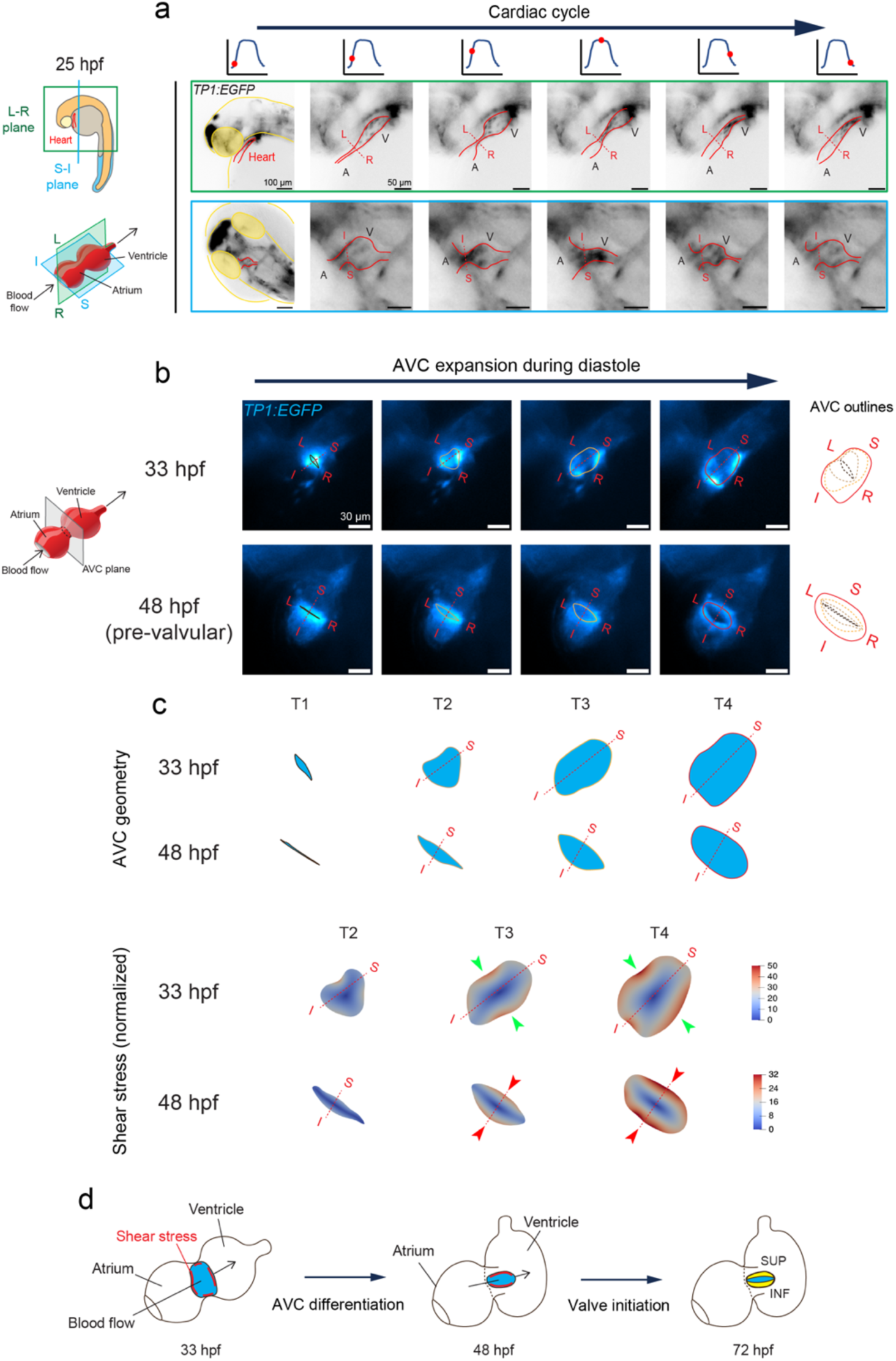
Peristaltic atrioventricular contraction aligns shear stress with the inferior-superior (S-I) axes during AV valve initiation. **(a)** At 25 hours post-fertilization (hpf), brightfield imaging of *Tg(TP1:EGFP)* embryos along two orthogonal planes of the heart tube (green and blue) throughout a cardiac cycle. Dashed lines indicate the corresponding valvular axes at the atrioventricular canal (AVC): superior-inferior (S-I) and left-right (L-R). See Supplementary Video 1. **(b)** At 33 and 48 hpf, brightfield imaging of *Tg*(*TP1:EGFP*) embryos along the AVC cross-section (gray plane) demonstrates the opening of the AVC along the superior-inferior (S-I) axis during diastole. By 48 hpf, the differentiated AVC becomes flattened along the S-I axis prior to the SUP/INF cusps. Colored lines delineate the AVC endocardial contours at various time points. See Supplementary Video 2. **(c)** Extracted atrioventricular canal (AVC) geometry from the brightfield images, and a 2D shear-stress simulation demonstrates the spatial variation in shear-stress along the L-R axis (green arrowheads) and the S-I axis (red arrowheads) at 48 hpf. The color scale indicates the normalized shear-stress for a steady, incompressible Poiseuille flow through the 2D AVC geometry. **(d)** Schematics showing the initial redistribution of shear stress before AV valve initiation. Anatomic labels: V, ventricle; A, atrium; AVC, atrioventricular canal; L-R, axis for LL and RL cusps; S-I, axis for SUP and INF cusps.

### Shear stress distribution aligns with valvular axes during 2- to 4-cusp transition

Before the initiation of SUP/INF cusps, the endocardial cells surrounding the atrioventricular canal (AVC) reduce their cytoplasmic volume and converge toward the center of the AVC, where elevated oscillation amplitudes of shear stress are observed. At 33 hours post-fertilization (hpf) and 48 hpf, the diastolic motion of the AVC endocardium is asymmetric and biased toward the S-I axis (**Figure 3b, Supplementary Video 2**). During AVC differentiation (36 - 48 hpf), its cross-section flattens along the S-I axis. Consequently, the shear stress simulation indicates a redistribution of high shear stress from the L-R axis to the S-I axis (**Figure 3c-d**).

At 14 dpf, mRNA staining of the shear stress-sensitive transcription factor Klf2a (gene: *klf2a*) and subsequent quantification (TP1-normalized) showed *klf2a* enrichment at the remodeled commissure (**Figure 4a-b**), consistent with the redistribution of a high-shear-stress region from the S-I axis to the L-R axis (**Figure 4c**). At 10 dpf, 4-D (time + space) light-sheet microscopy was performed to image *a Tg(TP1:EGFP)* larva whose LL, not RL, commissure has undergone remodeling (**Figure 4d, Supplementary Video 5**). From the extracted AVC geometry during a cardiac cycle, the 2D CFD simulation revealed a high-shear-stress region adjacent to the LL, not the RL, commissure (**Figure 4e**).

**Figure 4.**
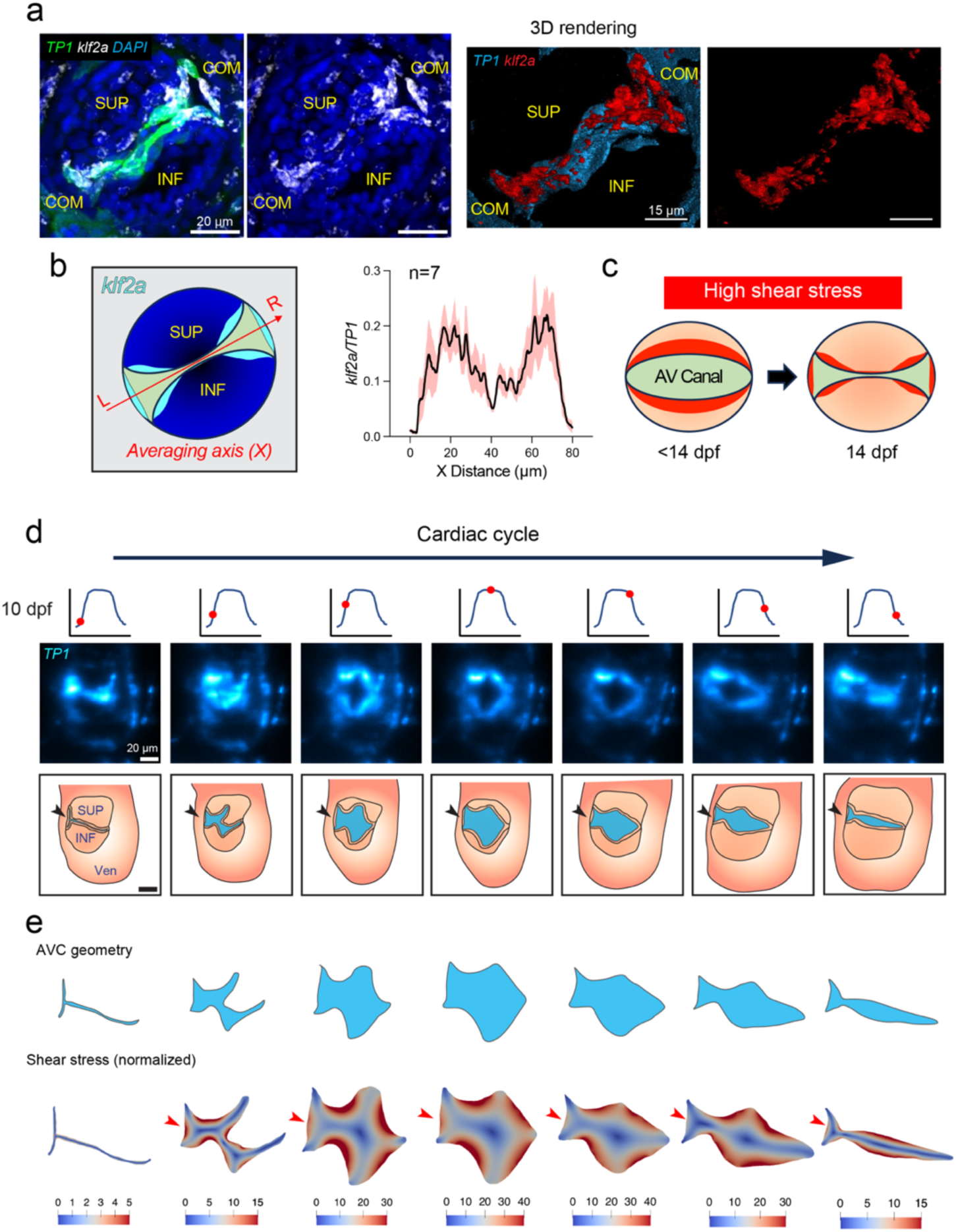
Increased endocardial wall shear stress occurs along the left-right lateral (L-R) axis during the 2- to 4-cusp transition. **(a)** At 14 dpf, whole-mount *in situ* hybridization for *klf2a* mRNA in the *Tg(TP1:EGFP)* heart reveals high *klf2a* expression around the remodeled commissures. The right panels show 3D reconstructions of Notch (*TP1*) and *klf2a* fluorescent signals. **(b)** Average valvular *klf2a* fluorescence intensity (*TP1*-normalized, black line) along the L-R axis peaks at the remodeled commissures (n = 7 hearts). The light-brown shading indicates the Standard Error of the Mean (SEM). **(c)** Schematics showing variations in the shear stress along the superior and inferior cusps and the L-R commissures during the 2-cusp to 4-cusp transition. **(d)** At 10 dpf, 4D light-sheet imaging of a *Tg(TP1:EGFP)* heart reveals the opening and closing of AV leaflets and the remodeling of the left lateral (LL) commissure over a cardiac cycle (arrowheads). See Supplementary Video 5. **(e)** AVC geometry extracted from the light-sheet images and 2D computational fluid dynamics (CFD) reveal a decrease in the shear stress along the remodeled LL commissure (arrowheads) but not along the right lateral (RL) commissure. The color scales indicate the normalized shear stress value under steady, incompressible Poiseuille flow through the 2D AVC geometry. Anatomic labels: SUP, superior cusp; INF, inferior cusp; COM, commissure; Ven, ventricle.

Hence, by 25 hpf, the S-I and L-R valvular axes are established by the asymmetric motion of the AVC endocardium and remain conserved relative to the ventricular orientation throughout cardiac morphogenesis. Prior to the initiation of SUP/INF (48 hpf) and LL/RL (14 dpf) cusps, the AVC remodels, and the shear stress distribution realigns with the corresponding valvular axis.

### Ventricle- vs. Atrium-driven shear stress to modulate 2- to 4-cusp transition

Multiple mechanosensitive molecular transducers, including Notch, Klf2a, and Ccm1 (or Krit1), coordinate in tandem to initiate the SUP/INF cusps ^26,29^. To test whether ventricular- and atrial-driven shear stress contributes differently to the initiation of AV valve leaflets via these genes, we investigate AVC formation utilizing two myosin heavy chain mutants, *haf* (“half-hearted”, *myh7^-/-^*) ^39^ and *wea* (“weak-atrium”, *myh6^-/-^*) ^40^.

In *haf* mutants, the mutation in ventricle-specific *myh7* disrupts ventricular contraction and results in incomplete atrial blood flow across the AVC into the ventricle (**Figure 5a, Supplementary Video 6**). At 6 dpf, whole-mount *in situ* hybridization for *notch1b*, *klf2a, and snai1b* (a mesenchymal marker for VICs ) mRNA in the WT heart (n = 7) shows clear colocalization in the SUP/INF cusps. In contrast, *haf* hearts (n = 7) show the absence of *notch1b* spatially converging toward the AVC and a complete loss of *snai1b*. In addition, four of seven hearts show no *klf2a* expression at the AVC (**Figure 5b**). As a result, *haf* mutants fail to generate unidirectional shear stress across the AVC to establish AVC polarity and valvular axes (**Figure 5c**). All *haf* mutants develop severe edema and die before the larval stage.

**Figure 5.**
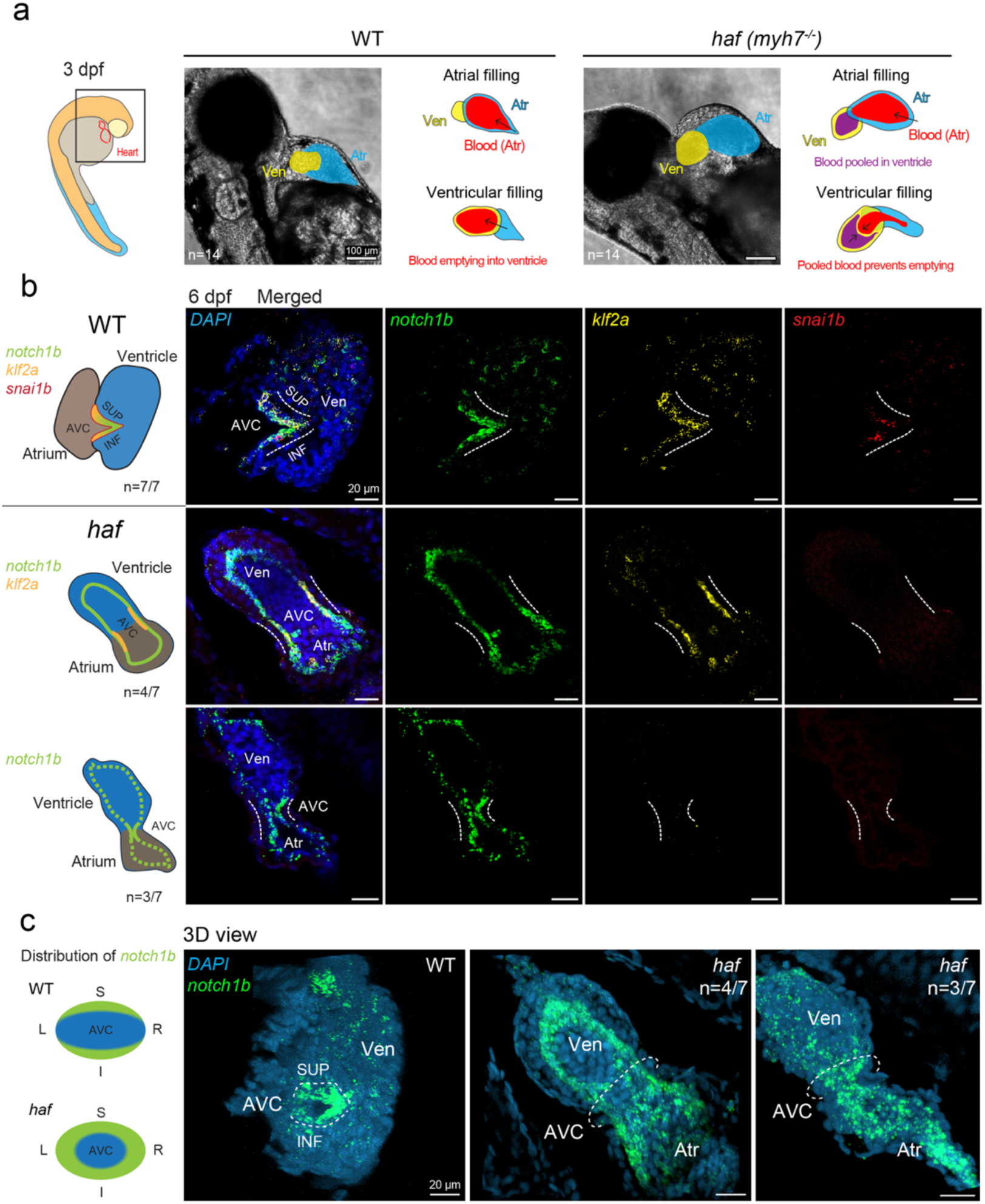
Ventricular contraction-generated shear stress across AVC is required to initiate the SUP/INF cusps. **(a)** At 3 dpf, brightfield imaging of *haf* (“half-hearted”, *myh7^-/-^*) mutants and their wild-type (WT) siblings. In WT hearts, the atrial blood (red) completely empties into the ventricle, whereas in *half* hearts, atrial inflow encounters resistance from the blood remaining in the non-contracting ventricle (purple), resulting in incomplete emptying and reduced flow across the AVC. See Supplementary Video 6. **(b)** At 6 dpf, whole-mount *in situ* hybridization for *notch1b*, *klf2a, and snai1b* mRNA in the WT heart (n = 7) shows clear colocalization in the SUP/INF cusps. In the *haf* hearts (n = 7), *notch1b* expression is no longer localized to the AVC, and *snai1b* expression is absent, suggesting the absence of ventricular interstitial cells (VICs). *klf2a* enrichment at the AVC occurs in four of seven hearts (4/7) but not in the remaining three (3/7). **(c)** 3D reconstruction of *notch1b* fluorescent signals reveals that *haf* mutants fail to develop peristaltic contractions along the left-right and inferior-superior axes. Anatomic labels: Ven, ventricle; Atr, atrium; AVC, atrioventricular canal; SUP, superior cusp; INF, inferior cusp.

In *wea* mutants, the atrial-specific *myh6* mutation disrupts atrial contraction and leads to a dilated atrium, where blood experiences prolonged stalling or flow reversal (regurgitation) (**Figure 6a, Supplementary Video 7**). In a cardiac cycle, the atrial filling phase in *wea* hearts is significantly shorter than in WT (n = 8, *p* < 0.0001 vs. *wea* = 6), whereas ventricular filling is similar (*p* = 0.0605) (**Figure 6b**). In our experience, approximately 59% of *wea* mutants survive to 28 dpf (81 out of 137). From 9 to 12 dpf, gross morphology and body length are not significantly different between *wea* mutants and their WT siblings, except that blood is pooled in the *wea* atrium (**Supplementary Figure 5a-b**).

**Figure 6.**
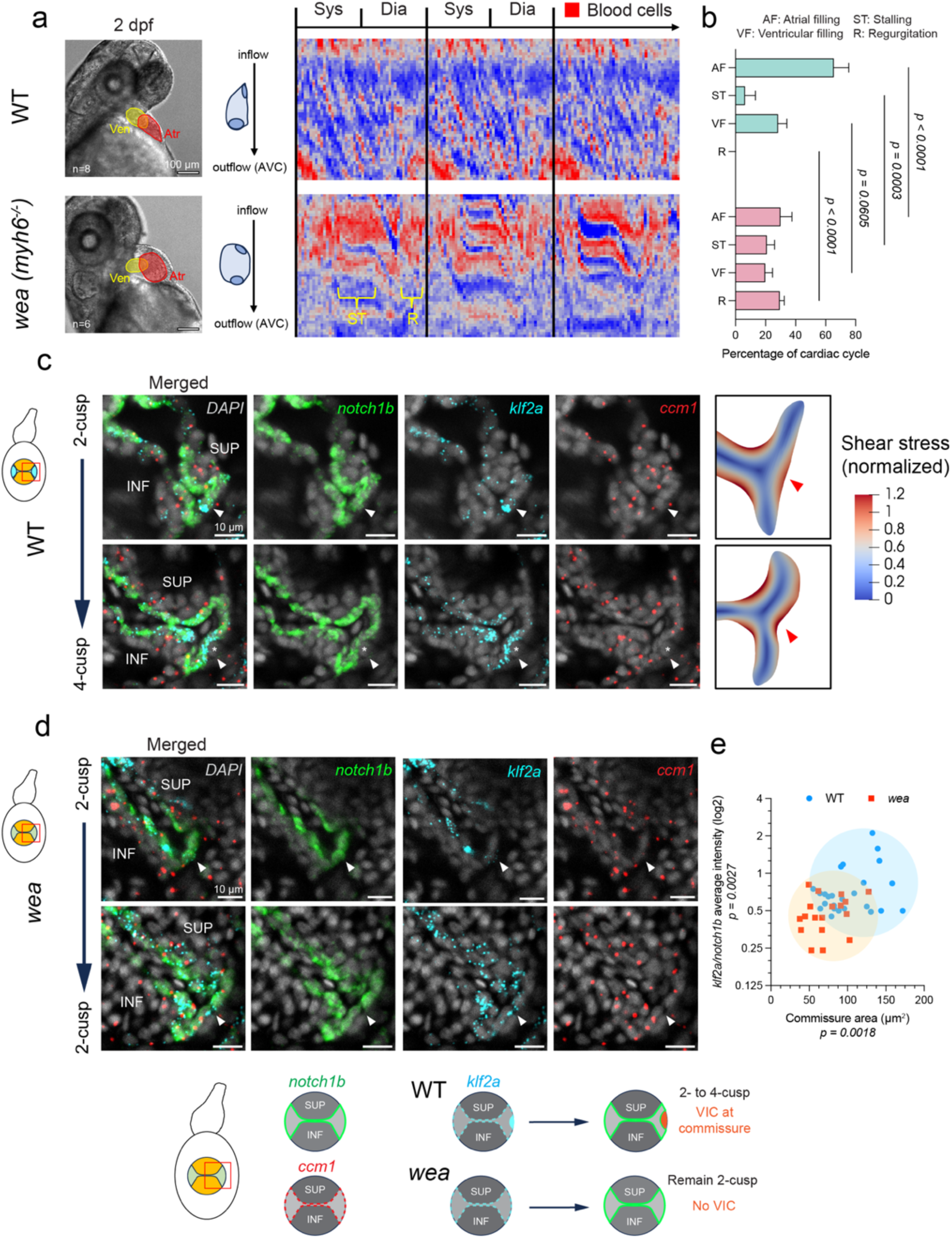
Atrial peristaltic contraction is required for 2- to 4-cusp transition. **(a)** At 2 dpf, brightfield imaging of *wea* (“weak-atrium”, *myh6^-/-^*) mutants and their wild-type (WT) siblings. The kymograph on the right shows blood flow dynamics across three cardiac cycles (Sys, systole; Dia, diastole). In WT hearts, blood flows across the AVC (top to bottom) during atrial contraction. In contrast, in *wea* hearts, blood experiences prolonged stalling (ST) or reversal/regurgitation (R) in the absence of atrial contraction. See Supplementary Video 7. **(b)** On average, *wea* mutants have significantly shorter atrial filling (AF) but longer stalling (ST) or regurgitation (R) periods than their WT siblings. Ventricular filling (VF) duration is similar between WT and *wea*. All values are reported as mean ± SD. Number of hearts analyzed: WT = 8, *wea* = 6. An ordinary two-way ANOVA followed by Sidak’s multiple-comparisons test was used to assess statistical significance. Source data are provided as a Source Data file. **(c)** At 14 dpf, whole-mount *in situ* hybridization for *notch1b*, *klf2a, and ccm1* mRNA in the WT heart (n = 15) shows enrichment of klf2a at the remodeled commissure (arrowheads) before (upper panels) and after VIC appearance (lower panels, asterisks). The spatial distribution of *klf2a* aligns with the spatial variations in the shear stress (right panels). **(d)** 86.7% of *wea* mutants in this study (26 of 30, compared with 1 of 38 WT) failed to undergo the 2-cusp-to-4-cusp transition, even though the remodeled commissure expanded (lower vs. upper panels). Enrichment of *klf2a* and VIC differentiation were not observed at the commissure (arrowheads). The spatial distribution of *notch1b* and *ccm1* was similar to that in WT hearts (quantification in Supplementary Figure 5). **(e)** Compared with WT, *wea* mutants developed significantly shorter commissures and lower average *klf2a* fluorescent intensity (*notch1b*-normalized) at the commissures. Each data point represents one commissure (n = 26 from 15 WT hearts, n = 18 from 12 *wea* hearts). Mann-Whitney test (for *klf2a/notch1b* intensity) and unpaired *t*-test (for commissure area) were applied to the means to determine statistical significance. Source data are provided as a Source Data file. Additional quantifications are shown in Supplementary Figure 5. Anatomic labels: Ven, ventricle; Atr, atrium; AVC, atrioventricular canal; SUP, superior cusp; INF, inferior cusp.

However, 86.7% of *wea* mutants in this study (26 of 30) fail to undergo the 2- to 4-cusp transition (i.e., no signs of VIC differentiation at either commissure), compared with 2.6% of WT (1 of 38). At 14 dpf, the spatial distribution of *klf2a* mRNA aligns with the shear stress around WT commissures (**Figure 6c**). *wea* hearts have significantly shorter commissures (*p* = 0.0018), fewer endocardial cells (*p* < 0.0001), and lower *klf2a* mRNA counts (raw measurement, *p* = 0.0215; *notch1b*-normalized, *p* = 0.0027) than WT hearts (n = 26 commissures from 15 WT larvae, n = 18 commissures from 12 *wea* larvae) (**Figure 6d-e and Supplementary Figure 5d**). By contrast, *notch1b* and *ccm1* mRNAs are evenly distributed along the AVC edge, and quantification at commissures is similar between *wea* and WT (**Supplementary Figure 5c**).

These findings suggest that ventricle-driven shear stress is required for establishing valvular axes and initiating SUP/INF cusps by regulating Notch and Klf2a. Atrium-driven shear stress is necessary for initiating the 2- to 4-cusp transition via Klf2a.

### Shear stress-activated VIC differentiation from endocardial cells during 2- to 4-cusp transition

At 14 dpf, the first VIC of the LL/RL cusps arises from endocardial cells at the remodeled commissure. The differentiating endocardial cell loses *notch1b* and *klf2a* but retains *ccm1* expression, consistent with the previously reported endothelial-mesenchymal transition (EndoMT) in SUP/INF cusps ^26,29^ (**Figure 7a**). It also loses endothelial identity (*fli1a:gal4*) and begins to express mesenchymal genes, including *snai1b*, which becomes highly enriched in later-cusp VICs (**Figure 7b-d**). The *wea* hearts contain a significantly lower percentage of *snai1b^+^*cells per commissure than the WT hearts (*p* < 0.0001, n = 21 commissures from 12 WT hearts, n = 17 commissures from 10 *wea* hearts), suggesting impaired VIC differentiation (**Figure 7e-f**). Staining for another VIC marker, *col1a2* , further demonstrates that *wea* mutants have significantly fewer total *col1a2^+^/snai1b^+^*VICs at the LL/RL commissure than the WT (*p* = 0.0002). The *col1a2* expression within SUP/INF cusps also shows a small, though significant, decrease in *wea* mutants (*p* = 0.0228, n = 11 WT hearts, 8 *wea* hearts) (**Figure 7g-h**). These results indicate that atrium-driven shear stress is critical for endocardial-VIC differentiation during the 2- to 4-cusp transition but less for the development of SUP/INF cusps.

**Figure 7.**
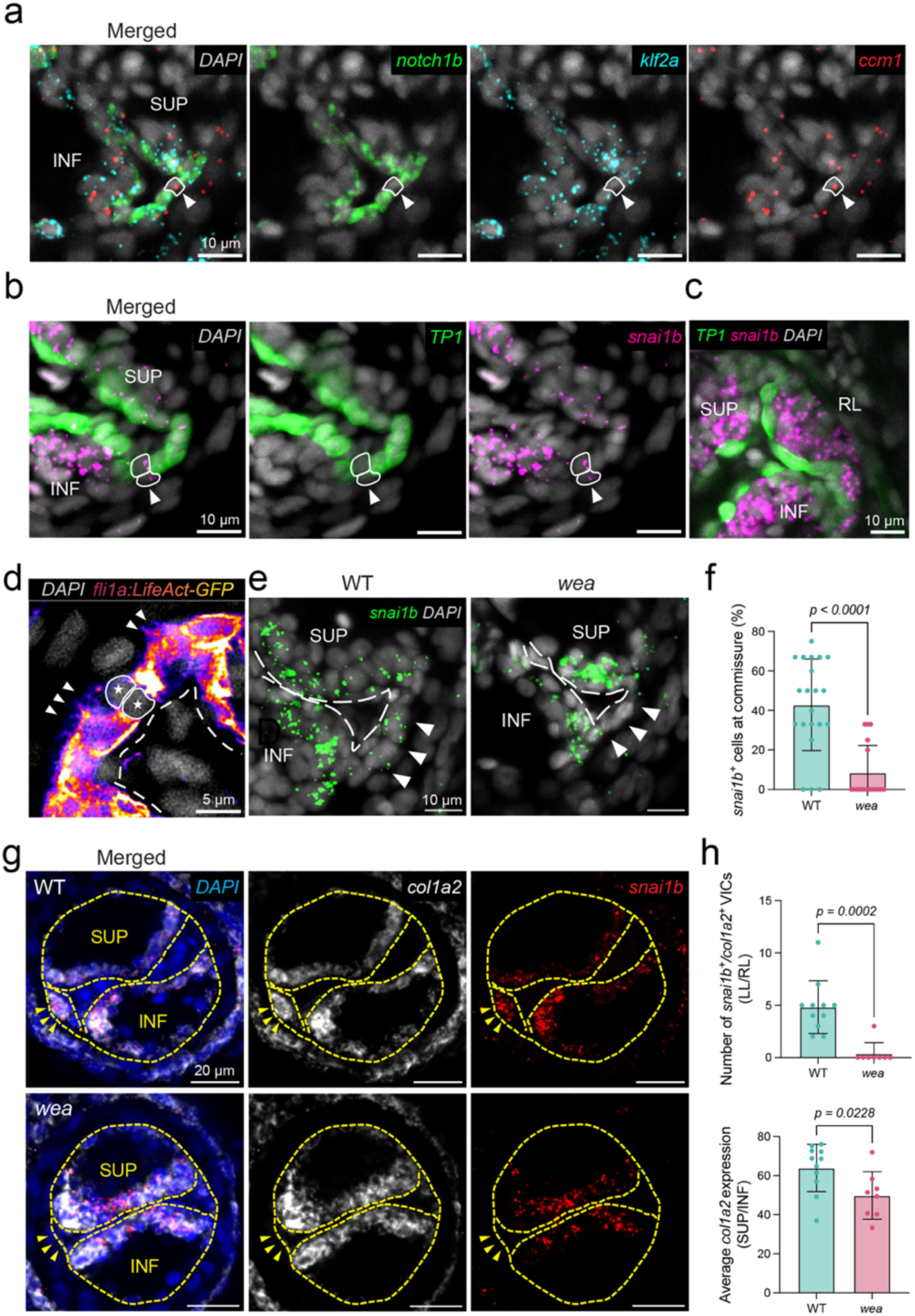
Shear stress at the remodeled commissure activates VIC differentiation. **(a)** The endocardial cell (solid outlines with arrowheads) undergoing differentiation loses *Notch1b* and *Klf2a* expression but retains *Ccm1* expression. **(b - c)** The Notch-negative (TP1) differentiating endocardial cells (solid outlines with arrowheads) express *snai1b* (mRNA), a mesenchymal gene involved in EndoMT. *snai1b* becomes highly enriched in VICs after the initiation of 4-cusps. **(d)** Endothelial identity (*fli1a:LifeAct-GFP* signals) is also lost during VIC differentiation (solid outlines with stars). Arrowheads mark endothelial filopodia. A dashed line outlines the free edge of the SUP/INF cusps and the remodeled commissure. **(e - f)** Compared with WT, *wea* mutants have significantly fewer *snai1b^+^*cells at the remodeled commissure (arrowheads). The dashed line outlines the free edge of the SUP/INF cusps and the remodeled commissure. All values are reported as mean ± standard deviation (SD). Each data point represents a single commissure (n = 21 from 12 WT hearts, n = 17 from 10 *wea* hearts). A Mann-Whitney test was applied to the means to assess statistical significance. Source data are provided as a Source Data file. **(g - h)** Compared with WT, *wea* mutants have significantly fewer *col1a2^+^*VICs (*snai1b^+^*) at the remodeled commissure (arrowheads). The average *col1a2* expression within the SUP/INF cusps also shows a small, though significant, decrease in *wea* mutants. The dashed line outlines the AVC and the free edge of the cusps. Number of hearts analyzed: n = 11 for WT, n = 8 for *wea*. All values are reported as mean ± SD. An unpaired *t*-test was used to assess statistical significance. Source data are provided as a Source Data file. Anatomic labels: SUP, superior cusp; INF, inferior cusp; RL, right lateral cusp.

### Spatial distributions of KLF4 and SNAI1 during the 2- to 4-cushion transition in the mouse AVC/OFT

Similar to zebrafish, from E10.5 to E11.5, the murine OFT and AV valves begin a 2-cushion-to-4-cushion transition ^42,43^, during which the two pre-existing cushions fuse and lateral (or intercalated for OFT) cushions emerge between them (**Figure 8a**). The two pairs of OFT and AV cushions later reorganize into the leaflets of the aortic/pulmonary and mitral/tricuspid valves, respectively. Compared with KLF2, KLF4 is more spatially and temporally specific in initiating EndoMT in mouse AV cushions. Snail family genes, *Snai1* and *Snai2* (*Slug*), are activated by TGF-β and Notch signaling, respectively, to regulate EndoMT ^19,44^. Because KLF2 lacks reliable antibodies, we stained for KLF4 and SNAI1 to examine their spatial distribution during the murine 2-cushion-to-4-cushion transition. Although the mouse AVC resembles a “dumbbell” shape, as in the zebrafish, KLF4 is distributed evenly within the AVC endothelium. It does not show spatial variation between cushions (**Figure 8b**). SNAI1 staining shows patterns akin to those in the zebrafish AVC during its transition, with highly enriched expression in the mesenchyme of pre-existing cushions and in differentiating endocardial cells at the commissure. Moreover, SNAI1^+^ cells exhibit weak or absent KLF4 expression, consistent with previous reports ^15^ and zebrafish studies. As the lateral AV cushion develops, SNAI1^+^ cells populate the bulk of it. On the other hand, the intercalated OFT cushions contain very few SNAI1^+^ cells, despite abundant KLF4 in the endocardium (**Figure 8c**). Therefore, KLF4 may not be the only mechano-sensitive transducer underlying the mouse 2-cushion-to-4-cushion transition.

**Figure 8.**
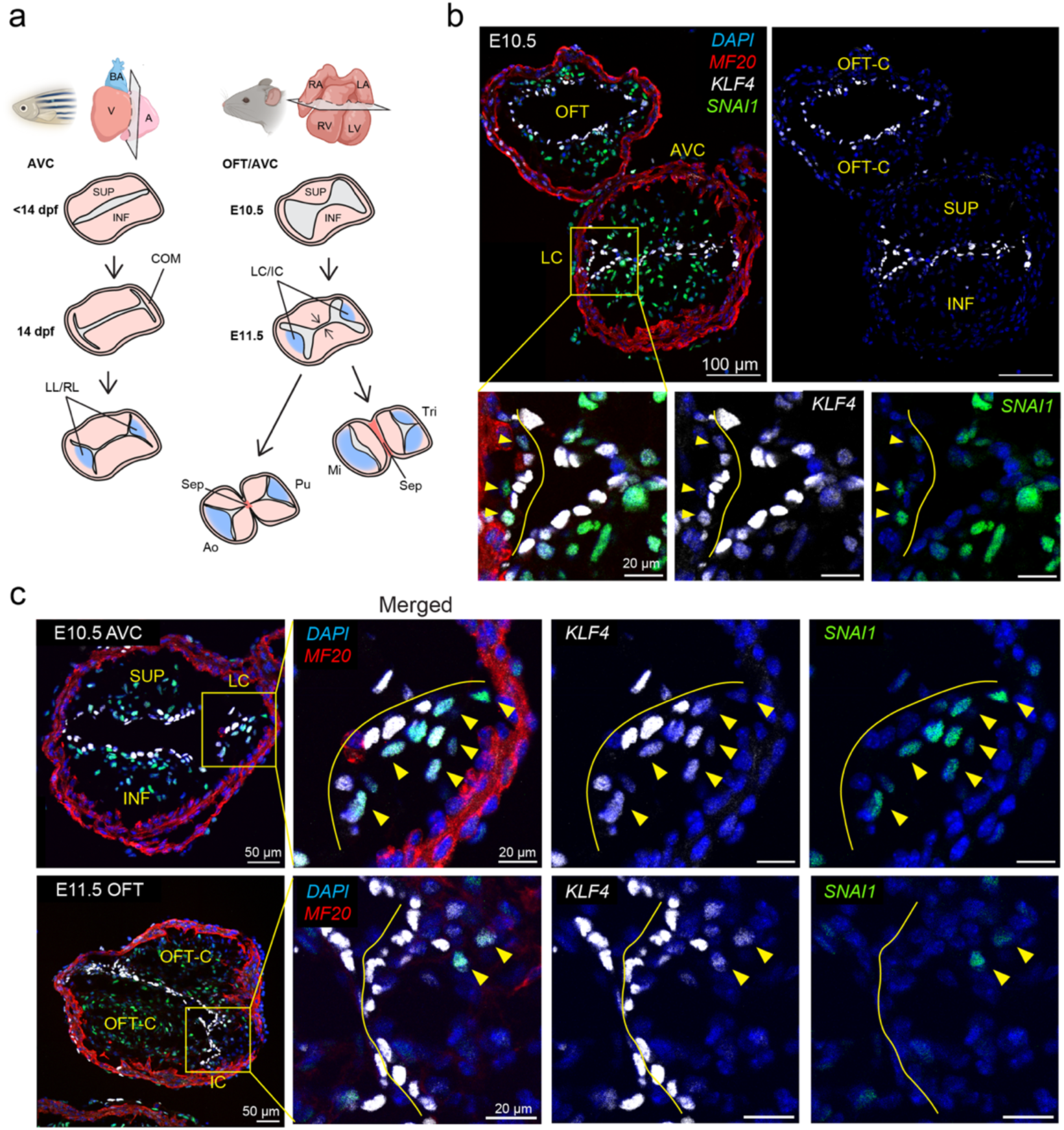
Spatial distributions of KLF4 and SNAI1 during the 2- to 4-cushion transition in the mouse AVC/OFT. **(a)** Schematic illustrating the 2- to 4-cushion transition during mouse AVC and OFT development. The SUP/INF cushions extend toward the center of the AVC/OFT and begin to fuse at E11.5. Meanwhile, the lateral (LC) and intercalated (IC) cushions develop along the orthogonal axis. The atrioventricular and OFT septa (Sep) arise from the fused cushions. The lateral and intercalated cushions become leaflets (blue shading) of the mitral/tricuspid (Mi/Tri) and aortic/pulmonary (Ao/Pu) valves, respectively. Partially created in BioRender. Wang, J. (2025) https://BioRender.com/jzz0scy **(b)** At E10.5, KLF4 and SNAI1 antibody staining shows colocalization at the initiating lateral cushion (solid outlines in the magnified lower panels). SNAI1^+^ cells (arrowheads) express weak KLF4 in the endocardium but not within the cushion, suggesting mesenchymal differentiation. The shape of KLF4^+^ endocardium resembles the dumbbell geometry, similar to the zebrafish AVC during the 2- to 4-cusp transition. **(c)** The intercalated cushion of OFT (solid outlines in the bottom panels) harbors very few SNAI1^+^ cells (arrowheads) compared with the lateral cushion of AVC (solid outlines in the top panels), suggesting a limited contribution from endocardial differentiation. Anatomic labels: AVC, atrioventricular canal; OFT, outflow tract; SUP, superior cusp; INF, inferior cusp; LL, left lateral cusp; RL, right lateral cusp; COM, commissure; LC, lateral cushions; IC, intercalated cushions; Sep, septum; Mi, mitral valve; Tri, tricuspid valve; Ao, aortic valve; Pu, pulmonary valve; OFT-C, OFT cushions.

## Discussion

Biomechanical signals orchestrate the formation of cardiac structures, which in turn shape the forces generated within the heart. Despite progress in understanding cardiac development, it remains unclear what governs the timing and location of growth of individual leaflets of the cardiac valve. Here, we present a novel mechanism underlying the patterning of valvular leaflets via chamber-specific shear stress. We uncover the role of atrial peristalsis (vs. ventricular syncytium) in the 2-cusp to 4-cusp transition of zebrafish atrioventricular (AV) valve. Employing two chamber-specific contractile mutants, 2-D shear stress analysis, and spatial transcriptomics, we demonstrate that the atrial shear stress aligning with the lateral valvular axis activates Klf2a-Snai1b signaling in Notch-negative/*ccm1*-positive endocardial cells to initiate bicuspid to quadricuspid valve transition. Thus, in the absence of atrial contraction (wea mutants), the 2- to 4-cusp transition (86.7%) fails to occur, and in the absence of ventricular contraction and relaxation (haf mutants), the formation of the first pair of leaflets (100%) fails to initiate.

Although scarce, evidence indicates that the acquisition of lateral cusps (or leaflets) in the AV valve is conserved among mammals, birds, and many ectothermic vertebrate . In our study, zebrafish and mouse AV valves exhibit considerable similarity prior to cardiac septation. Strikingly, zebrafish lateral cusps develop during the larval stage, whereas mammalian valves complete the transition during embryogenesis. To accommodate the growth of new cusps, the pre-existing leaflets separate from their original attachment points (i.e., commissures), thereby drastically reducing valvular cell proliferation. This mechanism allows the lateral cusps to reach 21.7% of the total AVC area by 45 dpf. It remains unclear whether this mechanism is conserved across species and what triggers remodeling of AV commissures at 14 dpf. EdU assays show that the separation involves the division of endocardial cells at the commissure, suggesting that specific signals delay their proliferation until the larval stage. These signals may also instruct the decline of VIC proliferation in SUP/INF cusps from 15 to 18 dpf and its recovery by 28 dpf. The good alignment between EdU data and cusp area measurements indicates that the main cellular event from 15 to 35 dpf is proliferation.

From 35 to 45 dpf, our RNA-seq data reveal downregulation of proliferation markers across the entire valvular tissue and upregulation of ECM remodeling genes, a pattern also observed in developing mouse AV valves. Among the several classes of upregulated genes, the chondrogenic pathway is the best recognized in valve development . We do not observe increased activation of *Sox9* (*sox9a*, *sox9b*), aggrecan (*acana*, *acanb*), and vimen in (*vim*), which have produced robust antibody staining in 1- and 2.5-year-old zebrafish AV valves ^47^. This indicates a temporal sequence of ECM secretion for valve maturation. Other components, including matrix metalloproteinase (*mmp2*), proteoglycan (*prg4*), glycoprotein (*pcolce2a*), collagen (*col2a1b*), and cytokeratin (*krt4*, *krt91*), are well characterized in mammalian valves and valvular diseases ^2,48,49^. Furthermore, we identified and validated a new valvular chondroitin sulfate proteoglycan, SNORC (Small NOvel Rich in Cartilage), recently discovered in human and mouse cartilages ^37^. Nevertheless, there is currently no functional study on the above genes in zebrafish valves.

Using a published spatial RNA sequencing dataset, we validated the expression of several unexpected genes. eEF1A1 (*eef1a1a*) is a member of translation elongation factors that regulate protein synthesis during the post-injury remyelination of the peripheral and central nervous systems ^50^. While eEF1A1 is predicted to be strictly embryonic in mammals, our data indicate that it is enriched in the AV valves of larval and adult zebrafish ^51,52^. Retinoic acid (RA) signaling plays an important role in cardiac development and in congenital heart disease ^53–56^. Retinol-binding protein 4 (RBP4, gene *rbp4*) is secreted by hepatocytes and adipose tissue to facilitate the transport of retinols. Clinically, increasing RBP4 mRNA in epicardial adipose tissue is associated with aortic stenosis ^57^. Here, we demonstrate that valves are an alternative source of RBP4. Similarly, human glycogenin-1 (GYG1, gene *gyg1b*) and liver glycogen phosphorylase (PYGL, gene *pygl*) are predominantly found in muscle (for glycogen synthesis) and the liver (for glycogen degradation), respectively ^58–60^. In zebrafish, both are highly expressed in the valves, especially *gyg1b*. Re-examining the expression of these genes in mammalian valves is needed, given their high level of similarity to zebrafish valves. Besides their developmental roles, it is also plausible to study their functions in zebrafish valve regeneration ^61–63^.

Understanding how chemical morphogens give rise to patterns and axes in the body through diffusion gradients and regulatory interactions has long been a scientific focus. However, recent studies demonstrate that the laterality of zebrafish hearts is established alongside the global body plan by flow-sensing cilia in the Kupffer’s vesicle ^64–66^. It is seldom appreciated that this laterality results in the early heart tube having a flat-elliptical cross-section ^67^. Our imaging data elucidate that the intrinsic axes of the heart tube are also the future valvular axes. Consistent with previous reports in zebrafish and chicken ^68,69^, we show that the collapsing motion of the AVC endocardium is asymmetrical and biased toward the S-I axis. Intriguingly, the long axis of the ellipse becomes the axis for the SUP/INF cusps after AVC differentiation, which flattens the AVC along the S-I axis and realigns the shear stress distribution. From there, the orientation and location of all cusps are determined. It is unclear how the direction of AVC flattening aligns with the S-I axis. Asymmetrical BMP signaling and ECM deposition around the AVC could both be underlying mechanisms ^70–72^.

What is more, we provide a more precise window (25 to 33 hpf) for the shift in ventricular contraction mode than some earlier accounts ^73,74^. Importantly, these time points precede the initiation of AVC differentiation (36 hpf) and cardiac looping (40 hpf), suggesting that the underlying mechanism may be upstream of these events. To the best of our knowledge, this study is the first to delineate the effects of ventricular vs. atrial shear stress on distinct pairs of valve leaflets. Although the AVC flow is believed to be predominantly atrial in zebrafish (E/A ratio < 1 during ventricular diastole) ^75,76^, loss of ventricular passive filling prevents the establishment of valvular axes. Some *haf* mutants show signs of narrowing or express *klf2a* in the AVC, but none exhibit proper notch1b *patterning*. In contrast, *notch1b* expression in *wea* mutants is unaffected. KLF4 staining in the mouse AVC and OFT outlines a geometry similar to that of the zebrafish AVC, which should produce a comparable shear stress distribution. However, we did not observe a clear spatial variation in KLF4 expression between the cushions. Therefore, further investigation will be needed to decipher the mechanical milieu at the AVC of these mutants and whether similar phenotypes can be expected in mammalian or avian hearts.

In mammalian hearts, the leaflets of the AVC and outflow tract (OFT) consist of cells from heterogeneous lineages. The four AV cushions are initially formed by endocardially derived *Tie2-Cre*^+^ cells, and the lateral cushions receive a large number of epicardially derived *Wt1-Cre*^+^ cells as the valve matures ^42,77^. The initial OFT cushions are endocardially derived before progenitors from the *Wnt1-Cre*^+^ neural-crest lineage invade the mesenchyme. Interestingly, the intercalated (lateral) cushions of OFT are formed by mostly *Tnnt2-Cre*^+^/*Isl1*^+^ second heart field (SHF) progenitors and receive minimal endocardial or neural-crest contributions ^43,78^. Similarly, the VICs are differentiated from SHF progenitors in the zebrafish OFT valve ^70^. The SUP/INF cusps of zebrafish AV valves are derived from the endocardium (*kdrl-Cre*^+^) and the neural crest (*sox10-Cre*^+^), without any epicardial contribution (*tcf21-Cre*^+^) ^27^. Our data shows that the earliest VICs of lateral AV cusps are derived from shear stress-activated endocardial cells. Extended lineage tracing experiments will elucidate whether epicardial progenitors join the VIC population in mature leaflets.

In summary, we provide a comprehensive characterization of the zebrafish AV valve’s 2-cusp to 4-cusp transition and uncover chamber-specific shear-stress patterning along valvular axes. Our study offers new perspectives on the origins of valvular structures and their interactions with biomechanical environments under hemodynamic shear stress.

## Methods

All animal experiments are conducted in accordance with protocols approved by the UCLA Institutional Animal Care and Use Committee (ID: ARC-2015-055 & ARC-2023-115).

### Zebrafish lines

Adult zebrafish are raised and bred in the UCLA Zebrafish Core Facility according to standard protocols ^79^. Embryos are cultured in E3 medium (5 mM NaCl, 0.17 mM KCl, 0.33 mM CaCl_2_, 0.33 mM MgSO_4_ in sterile diH_2_O) at 28.5 °C for all the procedures. 0.003% (w/v) 1-phenyl-2-thiourea (PTU, Sigma) is added to the medium to suppress the pigmentation. Embryos are transferred to the core facility at 6 days post-fertilization (dpf) and raised to 14 dpf or beyond. Transgenic line *Tg(flk:mCherry)* is provided by the UCLA Zebrafish Core. The *Tg(TP1:EGFP)^um14Tg^*line is a kind gift from David Traver at UCSD and Nathan Lawson at the University of Massachusetts Medical School ^35^. Myosin heavy chain mutants, *haf ^sk^*^24^ (“half-hearted”, *myh7^-/-^*) ^39^ and *wea ^m^*^58^ (“weak-atrium”, *myh6^-/-^*) ^40^, are a kind gift from Deborah Yelon at UCSD. *Tg(fli1a:Gal4)* and *Tg(UAS:LifeAct-GFP)* ^80^ are gifts from Julia Mack and Alvaro Sagasti at UCLA, respectively. Sex is not considered in our study design, as we focus on time points prior to sex differentiation in zebrafish (∼45 dpf).

### Mouse

Mice are housed in an animal facility under a 12-hour light/dark cycle at 22-24 °C and provided with standard irradiated mouse chow *ad libitum*. C57BL/6 mice (JAX, #000664) are bred for embryo generation and collection. The day of vaginal plug detection is designated as E0.5. Embryo sex is not considered a variable in our study.

### Whole-mount fluorescence in situ hybridization (FISH) and imaging

We have modified an established FISH-antibody staining protocol ^81^ according to the manufacturer’s recommendations for the RNAscope technology (Advanced Cell Diagnostics). Briefly, embryos are fixed in 4% PFA (paraformaldehyde) overnight at 4°C and washed in 0.1% PBST (Tween 20 in PBS). The skin around the heart is removed with fine forceps. For whole-heart imaging, larvae are euthanized by rapid cooling and fixed in 4% PFA overnight. After PBST washes, hearts are dissected. Embryos/hearts are treated with 100% methanol for at least 2 hours, then transferred to 3% (vol/vol) H_2_O_2_/methanol. After 1 hour of incubation, samples are rehydrated serially from 75% methanol (in 0.1% PBST) to 0.1% PBST. Tissue permeabilization is performed first with 1% Triton-X (in PBS, 1 hour) and then with RNAscope Protease Plus solution (30 minutes, 40°C). Next, RNAscope probes are added to the samples, and the FISH signal is developed step by step following the user manual for the RNAscope Multiplex Fluorescent v2 Assay. Opal 520, 570, and 690 (Akoya Biosciences, 1:1000) are used to develop the fluorescent signal for RNAscope probes. After FISH, anti-GFP (GeneTex GTX113617) primary antibody (1:100) is incubated with the samples in the co-detection antibody diluent (Advanced Cell Diagnostics) overnight at 4°C. Alexa Fluor™ 488 secondary antibodies (Invitrogen A-11008, 1:500) are then added along with DAPI to develop the signals for *TP1:EGFP* and for nuclear counterstaining. The hearts and embryos are mounted in 1% agarose on #1.5 cover glass (thickness ∼0.17 mm) for imaging.

Imaging is performed on a Leica SP8 scanning laser confocal microscope using the control software LAS X (version 3.5.7) from the Advanced Light Microscopy and Spectroscopy Lab at UCLA (RRID: SCR_022789). A 20X water-immersion lens (NA = 0.75, #506343) is used for the imaging. The probes and antibodies used in this study are listed in Supplementary Table 1.

### EdU proliferation assay

EdU (5-ethynyl-2′-deoxyuridine, Thermo Fisher) are diluted in the E3 medium to a concentration of 0.5 mM along with 0.5% dimethyl sulfoxide (DMSO) to label proliferating cells ^27^. Fish are incubated for 24 hours in 6-well plates (10 fish per well). After washing away the EdU, hearts are dissected and fixed. EdU labeling is detected using the Click-iT™ Plus EdU Cell Proliferation Kit for Imaging (Alexa Fluor™ 647 dye, Invitrogen).

### Immunofluorescence staining and imaging

Mouse embryonic hearts are isolated in chilled PBS and fixed in 4% PFA for 1 hour at 4 °C. After PBS washes, the hearts are saturated in 15% sucrose/PBS for 1 hour and in 30% sucrose overnight. Frozen blocks are prepared using the optimal cutting temperature (OCT) compound, with the hearts oriented for transverse sectioning at 10 µm. The frozen sections are washed in PBS and permeabilized in 0.25% Triton-X/PBS for 10 minutes. After 1 hour of blocking in 10% donkey serum (in 0.1% PBST), primary antibodies are added and incubated overnight at 4 °C. Finally, fluorescence is developed with a 1-hour secondary antibody incubation, and the slides are sealed in VECTASHIELD antifade mounting medium with DAPI (Vector Labs). Imaging is performed on a Leica SP8 scanning laser confocal microscope with a 20X (NA = 0.75, #506343) oil-immersion lens. The antibodies used in this study are listed in Supplementary Table 1.

### Widefield fluorescence imaging

To characterize the valvular axes during development, *Tg(TP1:EGFP)* embryos are anesthetized in 0.2 g/L tricaine solution (Sigma) and mounted in 1% low-melting-point agarose (Thermo Fisher). The embryos are imaged at each time point using an Olympus IX70 epi-fluorescent microscope (objective: UPLFLN 10X/0.3 NA) with a Prime BSI camera (exposure time: 10 ms, 2x2 binning, control software: Micro-Manager 1.4). Frames of minimal distance (between endocardium) are manually counted during the period in the atrium, ventricle, and AVC for statistical analysis. *Tg(fl1a:LifeAct-GFP), haf,* and *wea* embryos are imaged using the same method to capture their cardiac dynamics. To profile the hemodynamics in *wea* vs WT hearts, kymographs are constructed from the widefield videos using the Dynamic Reslice tool in FIJI ^82^. Care is taken to ensure that each embryo is imaged in the same orientation. Frames corresponding to atrial filling (AF), ventricular filling (VF), stalling (ST), and regurgitation (R) periods are manually counted for statistical analysis. Frames from three cardiac cycles are summed and divided by the total frames of three cycles to calculate the average percentage.

### Light-sheet fluorescence imaging

To visualize the motion of the transitioning AV valve at 10 dpf, *Tg(TP1:EGFP)* larvae are mounted in #1.5 glass-bottom dishes (Mattek) with 1% agarose and imaged using a Leica SP8 digital light-sheet (DLS) system (camera: Hamamatsu Orca Flash 4.0 V2; illumination objective: Leica HC PL FLUOTAR 2.5x/0.07 Dry; detection objective: Leica HC PL FLUOTAR 25x/0.95 Water) at 100 frames per second. The agarose around the gills and mouth is carefully removed to avoid suffocation. The larvae are anesthetized in 0.2 g/L tricaine (MS-222) solution (Sigma). At each z-position, 60 frames are acquired with a z-axis step size of 1 μm. The resulting 4D stacks are synchronized using a custom-written MATLAB program .

### Confocal image processing and quantification

To quantify the cusp areas, a maximum z-projection is first performed on the confocal z-stacks using FIJI . Then, ROIs are drawn around the individual cusps, and the areas of the superior and inferior (or left lateral and right lateral) cusps are averaged for each heart. The percentage of SUP/INF (or LL/LR) cusps in each heart is calculated as:

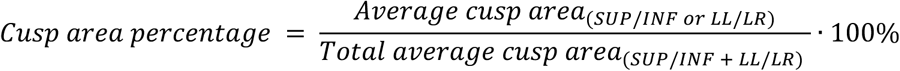

The percentage values from all hearts at each time point are then averaged for display.

The following logistic-growth equation is fitted to the average areas of SUP/INF (x) and LL/LR (Y) cusps in GraphPad Prism (v11.0.0):

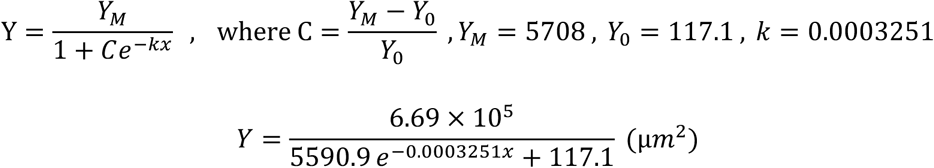

The inflection point (X_i_) indicates where the relative growth rate of lateral cusps reaches its maximum (i.e., where the second derivative of Y is zero). At this point, 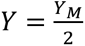:

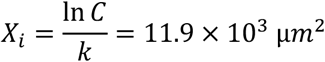

The point X_m_ marks the period of greatest increase in the relative growth rate, corresponding to the maximum of the second derivative:

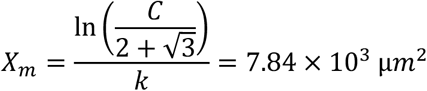

Counting of EdU^+^ cells was performed manually in FIJI or Leica LAS-X software across the entire 3D space of the cusps. The numbers in the superior and inferior (or left lateral and right lateral) cusps are averaged for each heart.

A custom Python script is executed in napari (v0.5.3) ^85^ to calculate *TP1*-normalized *klf2a* expression across the L-R axis of 14 dpf AV valves. First, pixels with TP1-EGFP fluorescence intensity > 10 are used as masks to isolate *klf2a* channel pixels at each Z-slice. Z-slices are selected to include the entire *TP1^+^* valvular region. Next, at each pixel along the L-R axis (across the entire image), klf2a and TP1 fluorescence intensities *a*re averaged over the orthogonal (S-I) axis and the Z axis. Data points outside the valvular region are manually removed. Finally, the resulting *klf2a* intensity is normalized by the *TP1* intensity and plotted (along with the standard deviation) in GraphPad Prism. The 3D rendering of *TP1* and *klf2a* fluorescent signals is also performed in napari’s native viewer.

To compare *notch1b* and *klf2a* expression between WT and *wea* hearts, ROIs are drawn to include all endocardial cells and VICs at the commissure. The average fluorescence intensity per pixel is measured for each channel before normalization. The expression of *col1a2* in SUP/INF cusps is measured in a similar fashion. The expression of *ccm1* is sparse, so the average number of mRNA molecules per cell is counted. For snorc, the total number of mRNA molecules per cusp is counted. Counting of *snai1b*^+^ and *snai1b^+^/col1a2^+^*cells is performed manually in FIJI or Leica LAS-X software across the entire 3D space of the commissure.

### 2-D shear stress simulation

Blood flow through the zebrafish atrioventricular canal (AVC) is modeled as steady, incompressible, laminar, two-dimensional (2D) Poiseuille flow. Contours of the AVC are manually traced from in vivo confocal images at multiple developmental stages using a custom interactive Python script. Unstructured triangular meshes are generated from these contours in Gmsh (v4.13.1) ^86^ with a prescribed mesh size. Meshes are exported in XDMF format for use with DOLFINx (v0.8.0), the finite element problem-solving environment of the FEniCS project ^87^.

Blood velocity through the 2D AVC contour is computed by solving the scalar Poisson problem derived from the steady-state Navier-Stokes equations (assuming Poiseuille flow):

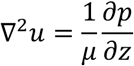

where 𝑢 denotes the axial (z) velocity component, 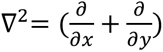 is the 2D Laplacian, 𝜇 is the dynamic viscosity, and 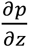 is the imposed pressure gradient across the valve. Homogeneous Dirichlet boundary conditions (𝑢 = 0) are imposed along the entire domain boundary to enforce the no-slip condition at the valve walls. The problem is discretized using first-order Lagrange finite elements and solved using a direct LU solver.

The shear stress magnitude is computed from the resulting velocity field as:

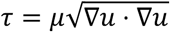

Because the analysis focused on spatial patterns of velocity and shear stress rather than absolute magnitudes, we set 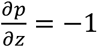 and 𝜇 = 1, yielding a non-dimensionalized formulation. Simulation outputs are then saved in VTU format and visualized using ParaView (v5.11.0) ^88^.

### Laser microdissection-assisted RNA sequencing

Hearts from 28-, 35-, and 45-dpf zebrafish are harvested in chilled PBS and embedded in OCT compound to produce frozen blocks. The atrium is removed, with the AV valve facing the cutting side during embedding. Cryosections of 16∼25 µm are mounted on polyethylene naphthalate (PEN) membrane slides (RNase- and DNase-free, Leica). Section thickness is optimized to ensure at least 3∼5 slices across the 3D space of the AV valve. Laser microdissection is performed on a Leica LMD7000 system. Dissected valve tissues are collected directly into the flat cap of a 0.2 mL PCR tube containing 65 µL of Buffer RLT (Qiagen) and stored at -80 degrees until further processing. 20 ng of carrier RNA (Qiagen) is added to the tissue lysate before freezing. In total, 30∼60 pieces (depending on age) of tissue cuttings (∼10 hearts) are pooled into each sample. Four biological replicates with no overlapping tissue sources are obtained for each time point (three from wild-type fish, one from *Tg(flk:mCherry; TP1:EGFP)*). RNA extraction is performed for all samples at once, following the RNeasy Micro Kit handbook (Qiagen).

Library construction (KAPA mRNA HyperPrep Kit) and RNA sequencing are performed on a NovaSeq 6000 platform (SP flow cell, Illumina) using paired-end 2 × 50 bp reads. Quality control (QC) is performed using NanoDrop (Thermo Fisher Scientific) and TapeStation (Agilent). Raw reads are downloaded from Illumina BaseSpace into UCLA Hoffman2 Cluster and quantified at the transcript level using Salmon (v1.10.2) ^89^ in selective alignment mode with a decoy-aware index. A combined transcriptome–genome (“gentrome”) reference is constructed from the Danio rerio assembly GRCz11 (release 110, Ensembl). The index is built using a k-mer size of 23. Quantification is performed with automatic library type detection, GC bias correction, and selective alignment.

The results are imported into R (v4.1.2) using tximeta (v1.12.4) ^90^ and summarized to gene-level counts. Differential expression analysis is performed using DESeq2 (v1.34.0) ^91^ to compare 35-and 45-dpf samples against 28-dpf samples as controls. Genes with low expression are filtered by retaining those with at least 10 counts in at least four samples. Principal component analysis (PCA) is performed on variance-stabilized counts using PCAtools (v2.6.0). One outlier from each time point (the *Tg(flk:mCherry; TP1:EGFP)* samples) is discarded due to low RNA quality in QC and distance from other samples in the PCA plot. Differential expression results are visualized using volcano plots generated with EnhancedVolcano (v1.12.0), with genes of interest highlighted. Up-regulated genes (*p.adj* < 0.05) from 35- or 45-dpf samples are first selected, then filtered based on the literature on valve biology. Additional valvular genes from the literature are added to the pool for examination. We employ this semi-supervised strategy to mitigate the effect of red blood cell contamination on traditional Gene Ontology (GO) analysis. The log-fold changes and Benjamini & Hochberg (1995) adjusted *p*-values are exported for visualization in GraphPad Prism. The original *p*-values are calculated from the Wald test in DESeq2. The final list of up-regulated genes is validated for spatial localization in the valves using a published Stereo-seq dataset of adult zebrafish hearts ^38^.

### Statistics and Reproducibility

With a few exceptions, all values are reported as mean ± standard deviation (SD). The statistical test used in each comparison is described in the corresponding figure legend. All statistical tests were performed in GraphPad Prism. A *p*-value less than 0.05 is deemed statistically significant. The resulting *p*-values and sample sizes (n) are provided in the corresponding figure legend. All data points for each statistical test were obtained from distinct samples. Details of each statistical analysis are provided in the corresponding sheet of the Source Data file.

All experiments in this study are repeated independently at least twice, yielding similar results. No statistical methods are used to predetermine the sample size. Sample sizes are determined based on published studies from our group, the availability of embryos/larvae, and the feasibility required to confirm the results. Overall, at least 5 samples per group (except for the RNA-seq analysis) are obtained for each statistical comparison.

In RNA-seq analysis, one outlier per time point is excluded due to low RNA quality in QC and distance from other samples in the PCA plot. When comparing the 2- to 4-cusp transition between WT and *wea* mutants, commissures with severely distorted morphology (based on staining) are not considered. No other exclusions are made. Embryos/larvae are randomly selected and distributed into experimental groups from a large pool of spawns. During imaging analysis, data are distributed to more than one investigator who did not perform the experiment, and they are blinded to the experimental conditions.

## Acknowledgments

We would like to thank the UCLA zebrafish core, David Traver (UCSD), Deborah Yelon (UCSD), Nathan Lawson (University of Massachusetts Medical School), Julia Mack and Alvaro Sagasti (UCLA) for the fish lines. We appreciate Hajime Fukui (Tokushima University) for advising the project. We are also grateful for the microscopy expertise from the UCLA Advanced Light Microscopy and Spectroscopy Lab (RRID: SCR_022789) and the RNA sequencing service from the UCLA Technology Center for Genomics & Bioinformatics (TCGB) (RRID: SCR_012204). This work used computational and storage services associated with the Hoffman2 Cluster which is operated by the UCLA Office of Advanced Research Computing’s Research Technology Group. This study is funded by NIH grants R01HL129727 (A.L.M., T.K.H.), R01HL159970 (A.L.M., T.K.H.), R01HL165318 (T.K.H.), and 2T32EB027629-06 (J.W., T.K.H.). A.D.K. was supported in part by NIH grant K25HL175208, AHA Career Development Award 24CDA1272816 and Stanford Maternal and Child Health Research Institute.

## Author contributions

Conceptualization, J.W. and T.K.H.; Methodology, J.W., A.L.B., P.Z., J.-N.C, B.Z., J.L., T.Y., S.-K.P., A.D.K., A.L.M., T.K.H.; Investigation, J.W., A.L.B., S.-K.P., C.Z.Z., and P.Z.; Writing – Original Draft, J.W., A.L.B., and T.K.H.; Writing – Review & Editing, J.W., A.L.B, P.Z., C.Z.Z., J.- N.C, B.Z., J.L., T.Y., S.-K.P., A.L.M., and T.K.H.; Funding Acquisition, A.L.M. and T.K.H.; Supervision, A.L.M. and T.K.H.

