## Supplementary Information for "A Continuum of Atrial Peristalsis Initiates the Bicuspid to Quadricuspid Valve Transition"

### Supplementary Figures and Table

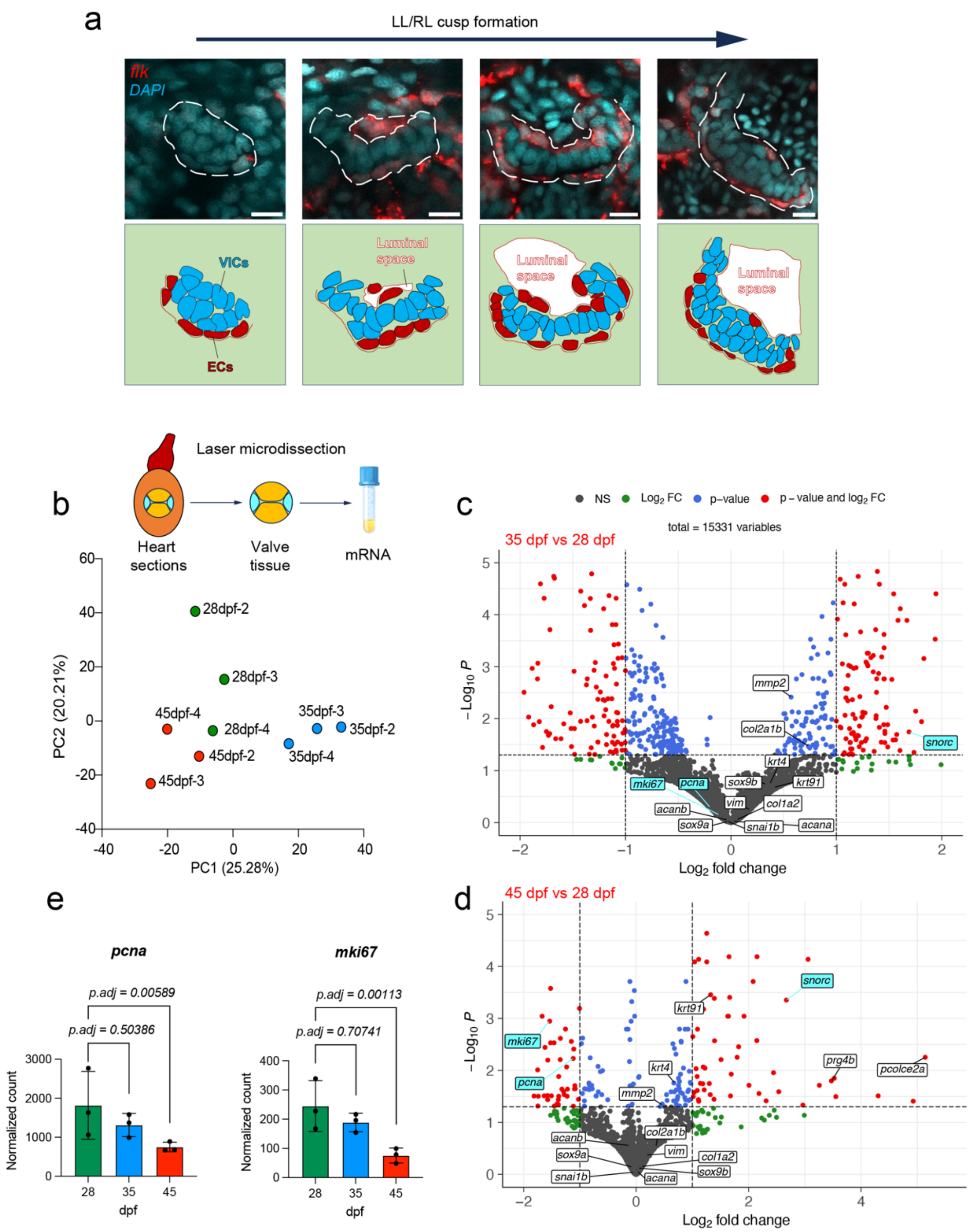
a
LL/RL cusp formation
flk
DAPI
VICs
ECs
Luminal space
Luminal space
Luminal space
b
Laser microdissection
Heart sections
Valve tissue
mRNA
28dpf-2
28dpf-3
28dpf-4
35dpf-2
35dpf-3
35dpf-4
45dpf-2
45dpf-3
45dpf-4
PC2 (20.21%)
PC1 (25.28%)
c
NS
Log2 FC
p-value
p – value and log2 FC
total = 15331 variables
35 dpf vs 28 dpf
Log2 fold change
d
45 dpf vs 28 dpf
Log2 fold change
e
pcna
p.adj = 0.00589
p.adj = 0.50386
Normalized count
dpf
mki67
p.adj = 0.00113
p.adj = 0.70741
Normalized count
dpf

**Supplementary Figure 1. Laser microdissection-assisted RNA sequencing of maturing AV valves. Related to Figure 1-2.**

**(a)** Confocal imaging of *Tg(flk:mCherry)* hearts demonstrates the formation of a lateral cusp after initiation at 14 dpf. Proliferating VICs form the initial cushion that protrudes into the AVC. The cushion then delaminates from the AVC wall to create the luminal space, and VICs form a monolayer cusp. The cusp continues to enlarge and mature with multilayer VICs.

**(b)** Laser microdissection-assisted RNA sequencing is performed on AV valve tissue from 28-, 35-, and 45-dpf WT zebrafish. Four samples from each time point are collected for sequencing, each containing 30~60 tissue pieces (depending on age) cut from ~10 hearts. One outlier from each time point is discarded based on the principal component (PC) analysis plot of normalized gene counts. Percentages of variance explained by the first two PCs are indicated on each axis. Source data are provided as a Source Data file.

**(c - d)** Volcano plots show the log-fold changes and Benjamini & Hochberg (1995) adjusted *p*-values for chondrogenic and mesenchymal genes of interest (labeled with lines) at 35 or 45 dpf versus 28 dpf. Two proliferation markers, *pcna* and *mki67*, as well as a novel chondrogenic gene, *snorc*, are highlighted in cyan. Original *p*-values are computed using the Wald test in DESeq2. Dashed lines mark the selected thresholds for log-fold changes (-1/1) and adjusted *p*-values (0.05). Colors indicate genes that pass the thresholds for log-fold changes (green), adjusted *p*-values (blue), or both (red). Normalized gene counts and adjusted *p*-values are shown in the subsequent bar charts.

**(e)** The normalized counts of proliferation markers *pcna* and *mki67* are significantly reduced in valvular tissues at 45 dpf compared with 28 dpf. Source data are provided as a Source Data file.

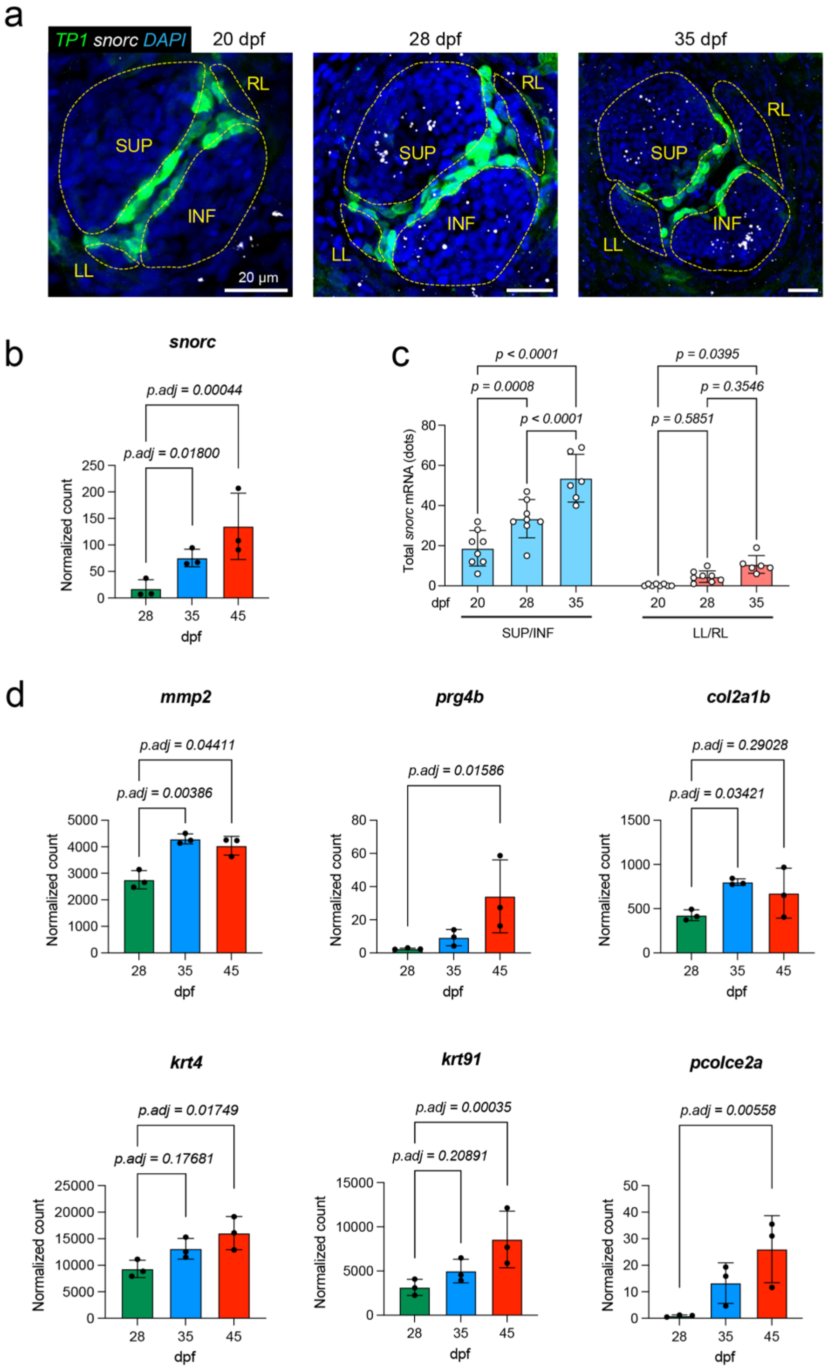
a
TP1 snorc DAPI
20 dpf
28 dpf
35 dpf
SUP
INF
RL
LL
20 µm
b
snorc
p.adj = 0.00044
p.adj = 0.01800
Normalized count
dpf
c
p < 0.0001
p = 0.0008
p < 0.0001
p = 0.0395
p = 0.3546
p = 0.5851
Total snorc mRNA (dots)
SUP/INF
LL/RL
d
mmp2
p.adj = 0.04411
p.adj = 0.00386
prg4b
p.adj = 0.01586
col2a1b
p.adj = 0.29028
p.adj = 0.03421
krt4
p.adj = 0.01749
p.adj = 0.17681
krt91
p.adj = 0.00035
p.adj = 0.20891
pcolce2a
p.adj = 0.00558

**Supplementary Figure 2. Chondrogenic genes in maturing AV valves. Related to Figure 2.**

**(a-c)** Whole-mount *in situ* hybridization of *snorc* mRNA in the *Tg(TP1:EGFP)* heart validates the mRNA sequencing results shown in panel (b). snorc expression i*ncreases* significantly in SUP/INF VICs from 20 to 35 dpf. In LL/RL cusps, expression does not differ until 35 dpf, indicating slower ECM maturation than in SUP/INF cusps. All values in panel (c) are shown as mean ± SD. Number of hearts analyzed: n = 8 for 20/28 dpf, n = 6 for 35 dpf. An ordinary two-way ANOVA followed by Sidak's multiple-comparison test was used to assess statistical significance. Source data are provided as a Source Data file.

**(d)** Normalized gene counts and Benjamini & Hochberg (1995)-adjusted *p*-values for chondrogenic genes that showed a statistically significant increase from 28 to 45 dpf. Source data are provided as a Source Data file.

Anatomic labels: SUP, superior cusp; INF, inferior cusp; LL, right lateral cusp; RL, right lateral cusp.

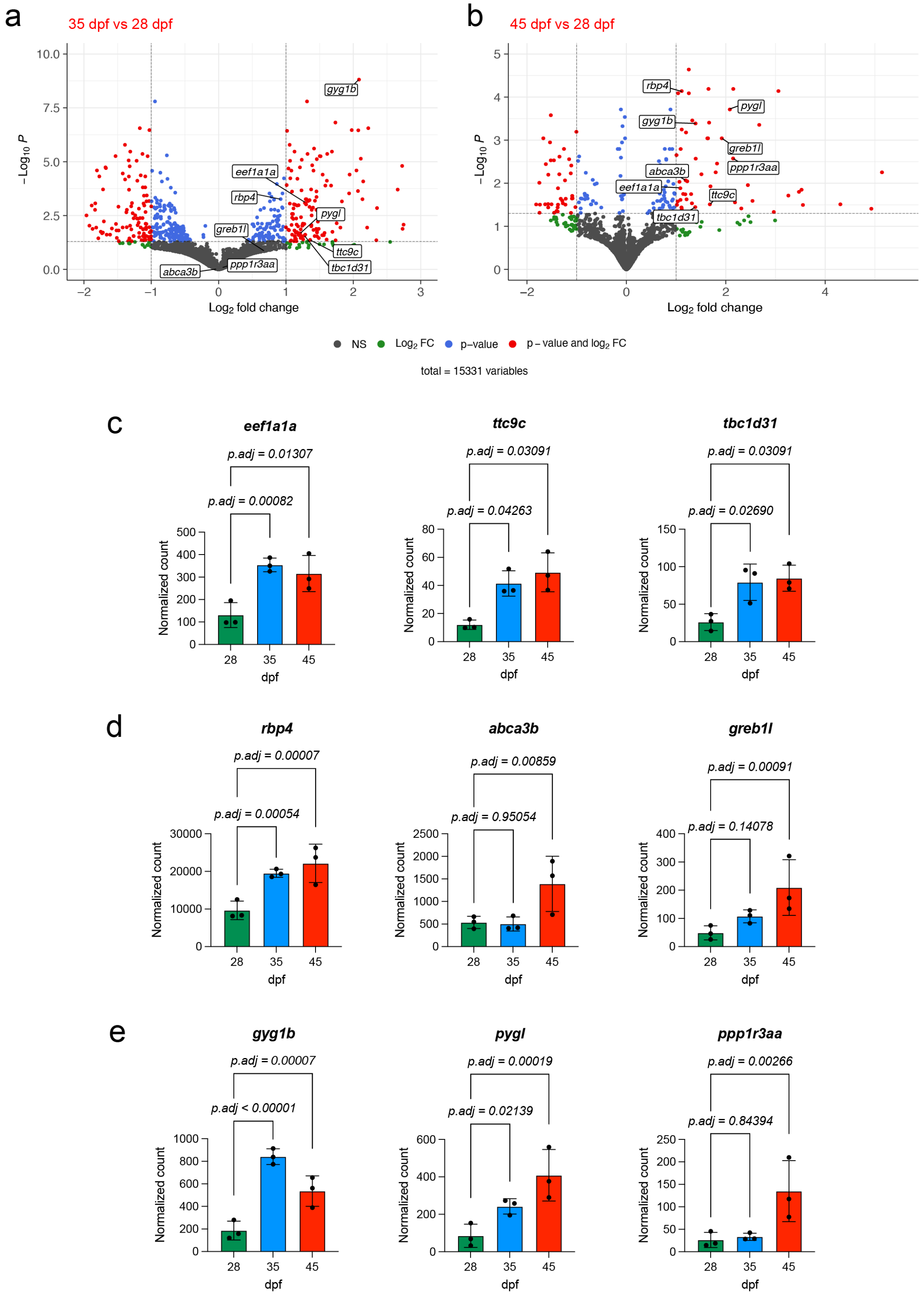
a
35 dpf vs 28 dpf
b
45 dpf vs 28 dpf
−Log10 P
Log2 fold change
gyg1b
eef1a1a
rbp4
pygl
greb1l
ttc9c
ppp1r3aa
tbc1d31
abca3b
NS
Log2 FC
p–value
p – value and log2 FC
total = 15331 variables
c
eef1a1a
p.adj = 0.01307
p.adj = 0.00082
ttc9c
p.adj = 0.03091
p.adj = 0.04263
tbc1d31
p.adj = 0.03091
p.adj = 0.02690
d
rbp4
p.adj = 0.00007
p.adj = 0.00054
abca3b
p.adj = 0.00859
p.adj = 0.95054
greb1l
p.adj = 0.00091
p.adj = 0.14078
e
gyg1b
p.adj = 0.00007
p.adj < 0.00001
pygl
p.adj = 0.00019
p.adj = 0.02139
ppp1r3aa
p.adj = 0.00266
p.adj = 0.84394
Normalized count
28
35
45
dpf

**Supplementary Figure 3. Genes involved in ciliation, retinoic acid signaling, and glycogen metabolism in maturing AV valves. Related to Figure 2.**

**(a-b)** Volcano plots show the log-fold changes and Benjamini & Hochberg (1995) adjusted *p*-values for genes related to protein synthesis, ciliation, retinoic acid signaling, and glycogen metabolism (labeled with lines) in AV valve tissue at 35 or 45 dpf versus 28 dpf. Original *p*-values are computed using the Wald test in DESeq2. Dashed lines mark the selected thresholds for log-fold changes (-1/1) and adjusted *p*-values (0.05). Colors indicate genes that pass the thresholds for log-fold changes (green), adjusted *p*-values (blue), or both (red). Normalized gene counts and adjusted *p*-values are shown in the subsequent bar charts.

**(c)** Normalized gene counts and Benjamini & Hochberg (1995)-adjusted *p*-values for protein synthesis- and ciliation-related genes that showed statistically significant increases from 28 to 45 dpf. Source data are provided as a Source Data file.

**(d)** Normalized gene counts and Benjamini & Hochberg (1995)-adjusted *p*-values for retinoic acid signaling-related genes that showed a statistically significant increase from 28 to 45 dpf. Source data are provided as a Source Data file.

**(e)** Normalized gene counts and Benjamini & Hochberg (1995)-adjusted *p*-values for glycogen metabolism-related genes that showed a statistically significant increase from 28 to 45 dpf. Source data are provided as a Source Data file.

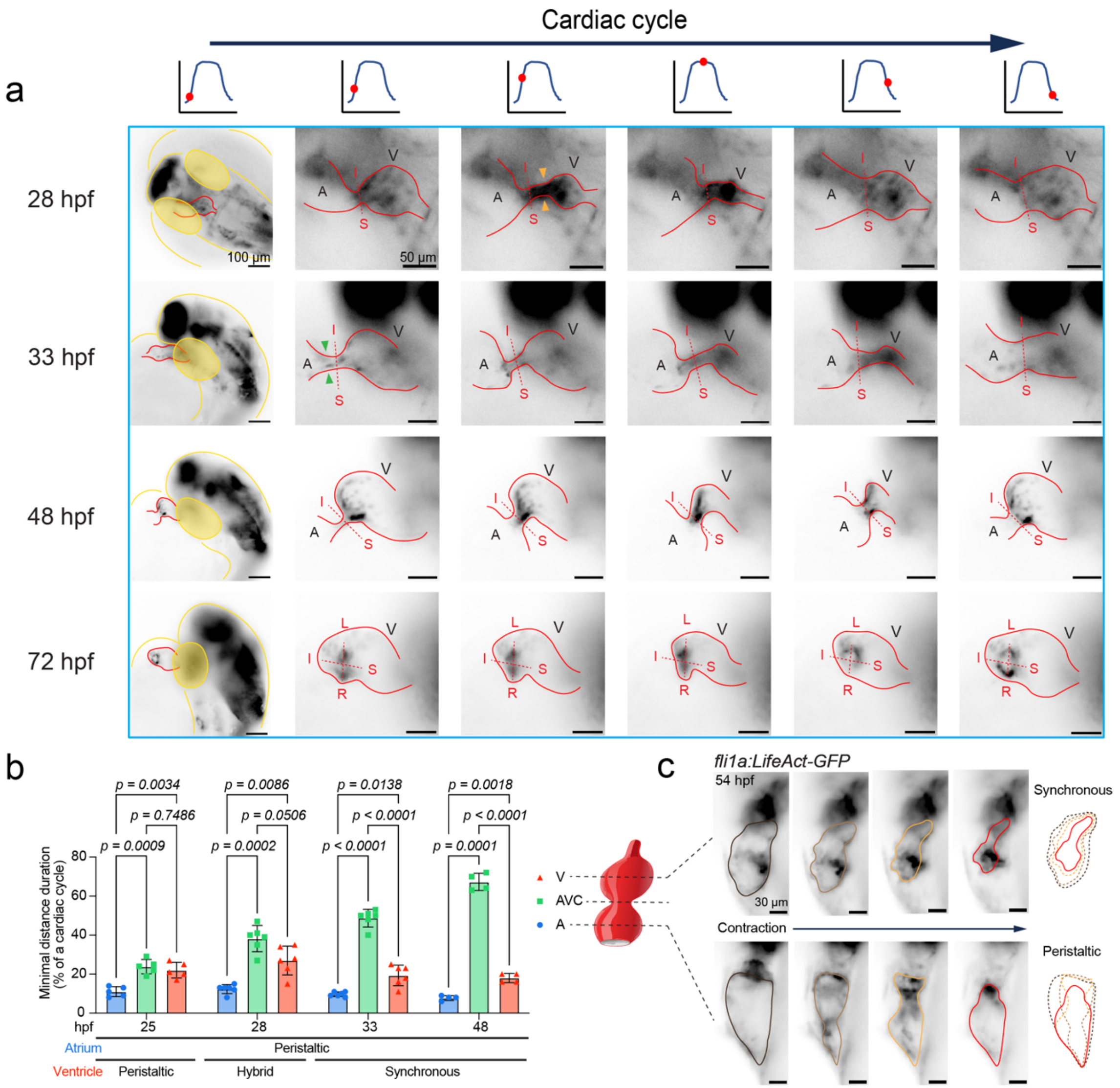


**Supplementary Figure 4. Peristaltic atrial-ventricular contraction followed by ventricular synchronous contraction during the initiation of AV valve. Related to Figure 3.**

**(a)** From 28 to 72 hpf, brightfield imaging of hearts of *Tg(TP1:EGFP)* Notch reporter embryos along the S-I axis (blue) demonstrates the formation of superior/inferior (SUP/INF) cusps. At 28 hpf, the peristaltic wave from the atrium reaches the mid-ventricle (orange arrowheads), while the rest of the ventricle contracts synchronously. At 33 hpf, the wave stops before the AVC (green arrowheads), and ventricular contraction becomes fully synchronous (i.e., a functional syncytium). See Supplementary Video 3.

**(b)** From 25 to 48 hours post-fertilization (hpf), peristaltic wave propagation was quantified by measuring the duration of endocardial wall contraction. The duration in the atrioventricular canal (AVC) and ventricle is significantly longer than in the atrium, indicating a slowing of the peristaltic wave at the AVC. At 33 hpf, the ventricular syncytium exhibits a substantially shorter duration than the AVC. All values are reported as means ± standard deviations (SD). *P*-values were assessed for each comparison. The number of *Tg(TP1:EGFP)* hearts analyzed was n = 7 at 25 hpf, n = 6 at 28/33 hpf, and n = 4 at 48 hpf. A mixed-effects model with the Greenhouse-Geisser correction, followed by Tukey's multiple-comparisons test on the means, was used to determine statistical significance. Source data are provided in the accompanying Source Data file.

**(c)** At 54 hpf, brightfield imaging of embryos expressing the endothelial actin reporter *Tg(fli1a:LifeAct-GFP)* reveals distinct peristaltic atrial and synchronous/syncytial ventricular contractions. Colored lines delineate the endocardial contour at different time points within one cardiac cycle. See Supplementary Video 4.

Anatomic labels: V, ventricle; A, atrium; AVC, atrioventricular canal; L-R, axis for LL and RL cusps; S-I, axis for SUP and INF cusps.

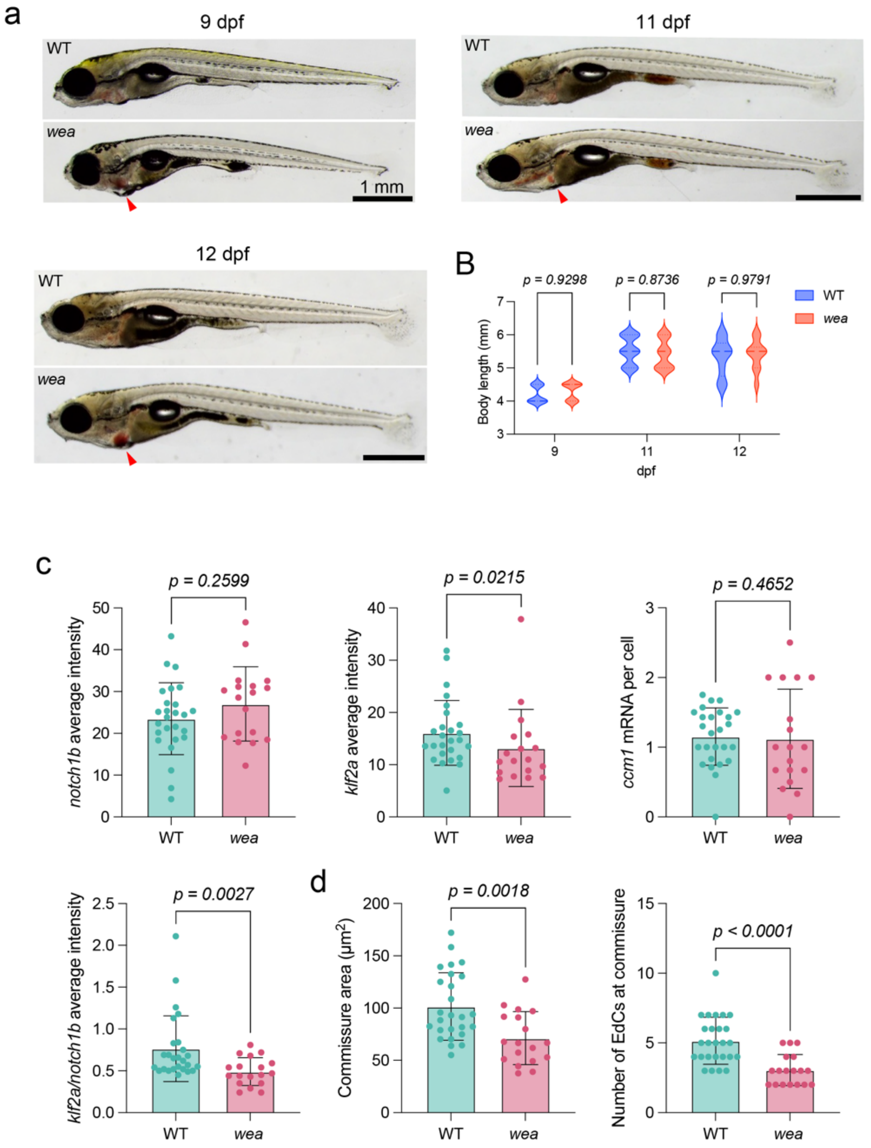


**Supplementary Figure 5. Morphological comparisons of *wea* mutants and WT siblings. Related to Figure 6.**

**(a-b)** The gross morphology and body length (violin plot) of *wea* mutants are comparable to those of wild-type (WT) from 9 to 12 days post-fertilization (dpf), except for pooled blood in the atrium (indicated by arrowheads). The width of each violin plot reflects the frequency of observations at specific body length values. The central line denotes the median, and the dotted lines represent the interquartile range (25th–75th percentile). The number of larvae analyzed includes: WT, n = 10 at 9 dpf, n = 17 at 11 dpf, and n = 9 at 12 dpf; *wea mutants*, n = 10 at 9 dpf, n = 14 at 11 dpf, and n = 10 at 12 dpf. An ordinary two-way ANOVA followed by Sidak's multiple-comparison test was used to assess statistical significance. Source data are provided in the accompanying Source Data file.

**(c)** At 14 days post-fertilization (dpf), expression levels of *notch1b* and *ccm1* at the commissure are comparable between *wea* and wild-type (WT) hearts. Conversely, klf2a *expression* and *the* normalized *notch1b-to-klf2a* ratio are markedly reduced at the commissure in *wea* hearts. All values are presented as means ± standard deviations (SD). Each data point represents measurements from one commissure (n = 26 for WT from 15 larvae; n = 18 for *wea* from 12 larvae). A Mann-Whitney test was used to assess the statistical significance of mean differences. The source data are available in a Source Data file.

**(d)** At 14 dpf, *wea* mutant hearts have a smaller commissure area and fewer endocardial cells (EdCs) at the commissure than WT hearts. All values are reported as mean ± SD. Each data point represents a measurement from one commissure (n = 26 WT from 15 larvae; n = 18 *wea* from 12 larvae). An unpaired t-test was used to assess statistical significance. Source data are provided as a Source Data file.

**Supplementary Video 1.**

Brightfield imaging of 25hpf *Tg(TP1:EGFP)* embryos along two orthogonal planes of the heart tube (green and blue) throughout a cardiac cycle. Related to Figure 3.

**Supplementary Video 2.**

At 33 and 48 hpf, brightfield imaging of *Tg*(*TP1:EGFP*) embryos along the AVC cross-section (gray plane) demonstrates the opening of the AVC along the superior-inferior (S-I) axis during diastole. Related to Figure 3.

**Supplementary Video 3.**

From 28 to 72 hpf, brightfield imaging of hearts of *Tg(TP1:EGFP)* Notch reporter embryos along the S-I axis (blue) demonstrates the formation of superior/inferior (SUP/INF) cusps. Related to Supplementary Figure 4.

**Supplementary Video 4.**

At 54 hpf, brightfield imaging of embryos expressing the endothelial actin reporter *Tg(fli1a:LifeAct-GFP)* reveals distinct peristaltic atrial and synchronous/syncytial ventricular contractions. Related to Supplementary Figure 4.

**Supplementary Video 5.**

At 10 dpf, 4D light-sheet imaging of a *Tg(TP1:EGFP)* heart reveals the opening and closing of AV leaflets and the remodeling of the left lateral (LL) commissure over a cardiac cycle (arrowheads). Related to Figure 4.

**Supplementary Video 6.**

At 3 dpf, brightfield imaging of *haf* ("half-hearted", *myh7*$^{-/-}$) mutants and their wild-type (WT) siblings. Related to Figure 5.

**Supplementary Video 7.**

At 2 dpf, brightfield imaging of *wea* ("weak-atrium", *myh6*$^{-/-}$) mutants and their wild-type (WT) siblings. Related to Figure 6.

#### Supplementary discussion

Across the animal kingdom, the design of cardiac valves generally reflects the species' physiological demands. For instance, the tricuspid OFT valve is a feature of endothermic hearts (e.g., mammals, birds), while most ectoderms, including fish, reptiles, and amphibians, have bicuspid OFT valves [1]. Multi-cuspid OFT valves can be found in the fast-swimming tuna (quadricuspid) and the largest known extant fish species, the whale shark (two rows of tricuspid), possibly an adaptation to the size of its heart [2]. Fluid-structure interaction simulation of the OFT valve in adult zebrafish predicts a low-pressure and laminar environment (mean Reynolds number = 1.25), without vortices or turbulence [3]. As the Reynolds number scales with size, one can expect hemodynamics to be more complex in the whale shark heart, underscoring the need for multiple leaflets to regulate flow. In chicken hearts, alteration of ventricular flow using a tungsten wire produces ventricular septal defects, extensive thickening of valvular tissue, and a lack of ECM stratification [5,6]. In mouse hearts, the removal of cardiac contraction (*Ncx1*$^{-/-}$) or primary cilia (*Ift20*$^{-/-}$) prevents EndoMT in AV cushions [7]. Mutations in the Notch pathway lead to the absence of EndoMT and VICs (*Notch1, RBPJk*), or BAV (*Sox17*) [8,9], while endocardial deletion of *Nfatc1* causes excessive EndoMT and impaired valve elongation [10]. Loss of KLF2/4-WNT signaling in valvular endocardium disrupts cushion remodeling and elongation, similar to the phenotypes in *Nfatc1* mutants [11]. Our study thus uncovered that spatial variation in the shear stress gradient patterns multi-cuspid valves.

### Supplementary Table 1

| REAGENT or RESOURCE | SOURCE | IDENTIFIER |
|---|---|---|
| Antibodies | | |
| Rabbit anti-GFP polyclonal | GeneTex | GTX113617 |
| Rabbit Snail (C15D3) monoclonal | Cell Signaling Technology | 3879 |
| Goat KLF4 polyclonal | R&D Systems | AF3158 |
| Mouse MF20 monoclonal | Invitrogen | 14-6503-82 |
| Goat anti-Rabbit IgG (H+L), Alexa Fluor 488 | Invitrogen | A-11008 |
| Goat anti-Rat IgG (H+L), Alexa Fluor 594 | Invitrogen | A-11007 |
| Goat anti-Mouse IgG (H+L), Alexa Fluor 647 | Invitrogen | A-21235 |
| Chemicals, Peptides, and Recombinant Proteins | | |
| 1-phenyl-2-thiourea (PTU) | Sigma-Aldrich | P7629 |
| Tricaine/MS-222 | Sigma-Aldrich | E10521 |
| TopVision Low Melting Point Agarose | Thermo Fisher | R0801 |
| DAPI | Thermo Fisher | D1306 |
| EdU (5-ethynyl-2′-deoxyuridine) | Thermo Fisher | A10044 |
| VECTASHIELD antifade mounting medium with DAPI | Vector Labs | H-1200-10 |
| PEN membrane (2 µm) coated glass slides, RNAse free | VWR (Leica 11505189) | 76414-898 |
| RNAscope Protease Plus | Advanced Cell Diagnostics | 322331 |
| Critical Commercial Assays | | |
| RNAscope™ Multiplex Fluorescent Detection Kit v2 | Advanced Cell Diagnostics | 323110 |
| RNAscope Probe-Dr-snai1b | Advanced Cell Diagnostics | 505101-C3 |
| RNAscope Probe-Dr-col1a2 | Advanced Cell Diagnostics | 526061-C2 |
| RNAscope Probe-Dr-notch1b | Advanced Cell Diagnostics | 431941 (C1) |
| RNAscope Probe-Dr-klf2a | Advanced Cell Diagnostics | 504421-C2 |
| RNAscope Probe-Dr-snorc | Advanced Cell Diagnostics | 1588411-C2 |
| RNAscope Probe-Dr-krit1 (ccm1) | Advanced Cell Diagnostics | 1794011-C3 |
| Opal 520 | Akoya Biosciences | NC1601877 |
| Opal 570 | Akoya Biosciences | NC1601878 |
| Opal 690 | Akoya Biosciences | NC1605064 |
| Co-Detection Antibody Diluent | Advanced Cell Diagnostics | 323160 |
| Click-iT™ Plus Alexa Fluor™ 647 Picolyl Azide Toolkit | Thermo Fisher | C10643 |
| RNeasy Micro Kit (50) | Qiagen | 74004 |
| Experimental Models: Organisms/Strains | | |
| *haf* $^{sk24}$ *(“half-hearted”, myh7* $^{-/-}$*) and wea* $^{m58}$ *(“weak-atrium”, myh6* $^{-/-}$*)* | Yelon Lab and Hsiai Lab | *Auman et al.*, PLoS Biol (2007), *Berdougo et al.*, Development (2003) |

| | | |
|---|---|---|
| *Tg(flk:mCherry)* | UCLA fish core and Hsiai Lab | N/A |
| *Tg(fli1a:Gal4) and Tg(UAS:LifeAct-GFP)* | Mack Lab, Sagasti Lab, and Hsiai Lab | *Helker et al.,* Development (2013) |
| *Tg(TP1:EGFP)* | Traver Lab, Lawson Lab, and Hsiai Lab | *Parsons et al.*, Mechanisms of Development (2009) |
| *C57BL/6 mice* | JAX | 000664 |
| Software and Algorithms | | |
| FIJI (v2.9.0) | Schindelin *et al.*, Nature Methods (2012) | RRID:SCR_002285 |
| MATLAB (R2019a) | MathWorks | RRID:SCR_001622 |
| GraphPad Prism (v9.5.0) | GraphPad Software | RRID:SCR_002798 |
| ParaView (v5.11.0) | Kitware | RRID:SCR_002516 |
| Python (v3.12.6) | Python Software Foundation | RRID:SCR_008394 |
| napari (v0.5.3) | conda-forge | RRID:SCR_022765 |
| Gmsh (v4.13.1) | Python | RRID:SCR_021226 |
| DOLFINx (v0.8.0) | Python | N/A |
| Salmon (v1.10.2) | bioconda | RRID:SCR_017036 |
| R (v4.1.2) | R Core Team | RRID:SCR_001905 |
| RStudio (v2023.03.0+386) | Posit Software | RRID:SCR_000432 |
| tximeta (v1.12.4) | Bioconductor | RRID:SCR_028005 |
| DESeq2 (v1.34.0) | Bioconductor | RRID:SCR_028005 |
| PCAtools (v2.6.0) | Bioconductor | RRID:SCR_025593 |
| EnhancedVolcano (v1.12.0) | Bioconductor | RRID:SCR_018931 |
